# Systematic Discovery of Pathogen Effector Functions across Human Pathogens and Pathways

**DOI:** 10.1101/2025.11.17.687821

**Authors:** Tomas Pachano, He Leng, Guillaume Dugied, Travis Tribble, Vincent Loubiere, Felix Rauh, Yeojin Lee, Alexander Schleiffer, Veronika Young, Benjamin Weller, Eleanor A. Lyons, Matthew R. Hass, Leah C. Kottayan, Matthew T. Weirauch, Juan I. Fuxman Bass, Hayley J. Newton, Alexander W. Ensminger, Pascal Falter-Braun, Daniel Schramek, Alexander Stark, Mikko Taipale

## Abstract

Pathogens deploy effector proteins to exploit host cell biology, and most pathogen open reading frames (ORFs) are rapidly evolving and lack functional annotation. We developed the eORFeome, a scalable functional genomics platform encompassing 3,835 effector ORFs from diverse viruses, bacteria, and parasites. High-throughput barcoded screens across NFκB, apoptosis, p53, cGAS–STING and MHC-I pathways revealed functions for hundreds of uncharacterized eORFs, unexpected new activities for known effectors, and distinct pathway-specific functions encoded by single ORFs. Illustrating the power of the approach, we identify HHV6A U14 as a p53 antagonist, HHV7 U21 as a dual-function STING antagonist and MHC-I antigen display inhibitor, and adenoviral 13.6K/i-leader protein as a *de novo* evolved TAP inhibitor that suppresses MHC-I display. These results establish a general framework for systematic effector annotation, uncover new mechanisms of host–pathogen interaction across kingdoms, and highlight pathogen effectors as a versatile toolkit for rewiring and probing human cellular pathways.

## INTRODUCTION

Pathogens express specialized gene products, known as effectors, to manipulate host molecular processes and establish conditions that are favorable for their survival and transmission.^1–3^ Many bacteria and eukaryotic parasites have evolved secretion systems that inject effector proteins into the host cell during infection, whereas viral effectors are mostly produced by the host ribosomes after viral entry (**Figure 1A**). By interfacing directly with host cellular machinery, effector proteins modulate immune responses, alter signaling pathways, and hijack metabolic processes to create a conducive environment for the pathogen.

**Figure 1.**
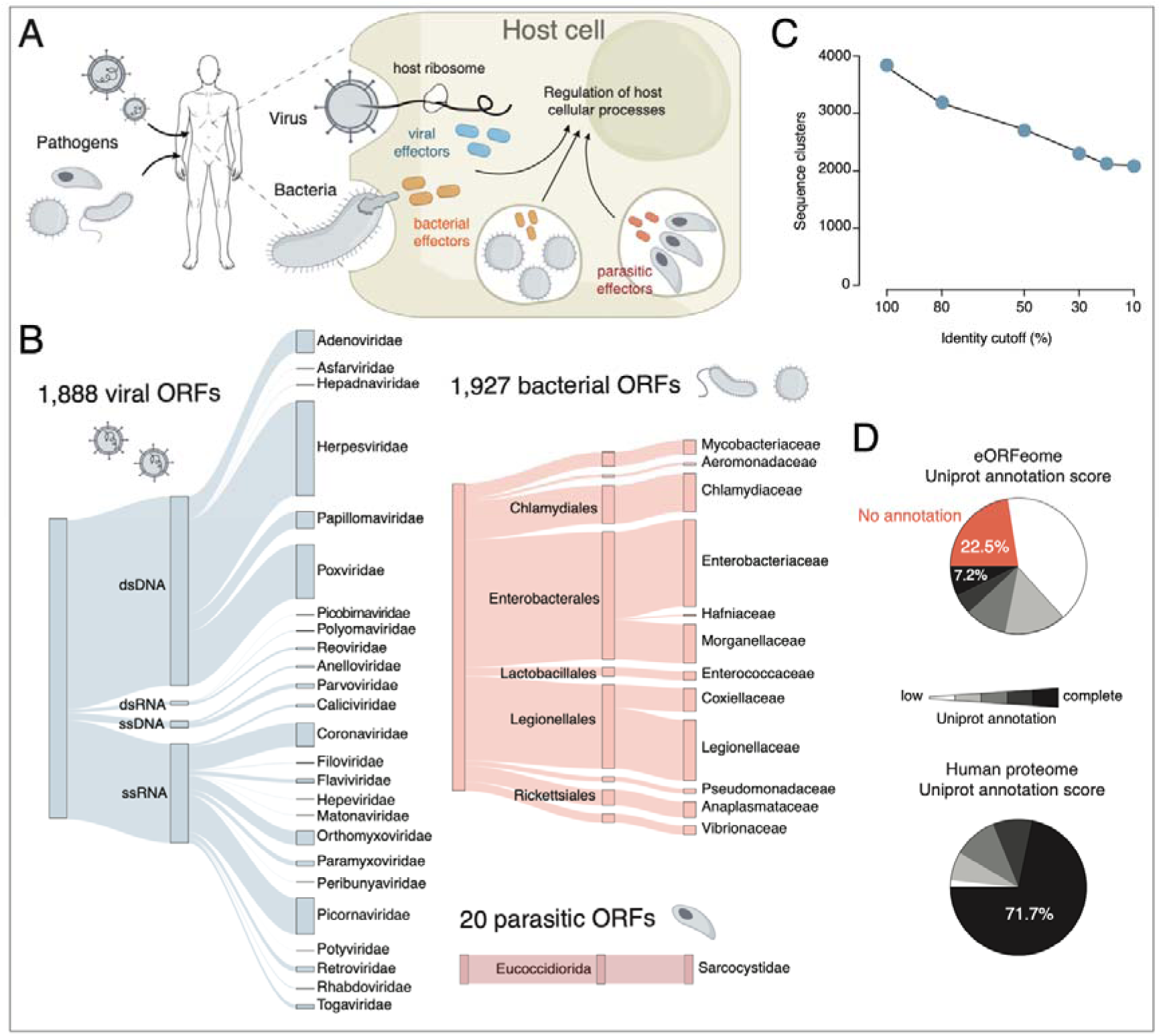
The Pan-Pathogen eORFeome Library. (A) Schematic of host cell manipulation by pathogen effectors. Diverse pathogens, including viruses, bacteria, and parasites, deploy effector proteins into host cells to hijack cellular processes, thereby promoting pathogen survival and subverting immune responses. (B) Alluvial diagrams illustrate the phylogenetic representation of the viral, bacterial and parasitic eORFs present in the eORFeome library. The flow diagrams connect taxonomic ranks - from Baltimore classification to Family for viruses, and from Order to Family for bacteria and parasites- with stream width corresponding to the abundance of ORFs from each group. (C) Sequence homology analysis of the eORFeome library. eORFs were clustered using MMseqs2^106^ at varying sequence identity cutoffs. The plot shows the total number of resulting homology clusters for each identity threshold. (D) Characterization of the eORFeome library (top) or human proteome (bottom) by UniProt annotation score. The pie chart shows the percentage of ORFs corresponding to each score. The grayscale colour key indicates the annotation level, from score 1 (white) to score of 5 (black). Orange corresponds to no entry in the UniProt database.

At least 2,000 pathogenic viruses, bacteria, and parasites are known to infect humans^4–6^ and the increasing availability of pathogen genome sequences has facilitated the identification of millions of viral open-reading frames (ORFs)^7,8^ and the computational prediction of thousands of bacterial effectors.^9–12^ However, with the exception of a few well-studied bacteria and viruses,^13–17^ the vast majority of pathogen effectors remains uncharacterized. This leaves a large reservoir of pathogen proteins that are highly enriched for novel functions shaped by the unremitting arms race between pathogens and their hosts. Indeed, most viral proteins lack structural homologs in the AlphaFold database,^18^ pointing to unique functions and mechanisms of action. Similarly, studies of secreted bacterial effectors have revealed many unexpected molecular mechanisms, including unique post-translational modifications.^19^

The fact that thousands of sequenced pathogen ORFs have not been characterized has far-reaching consequences for many fields, both scientific and societal. Pathogenic effector proteins target functions and pathways relevant to cell biology and human health, particularly to infection, inflammation, and cancer.^20–23^ Systematic functional characterization of these effectors could uncover novel host vulnerabilities and reveal potential therapeutic entry points, thus pre-emptively illuminating mechanisms by which emerging pathogens might manipulate human cells. Moreover, the effectors constitute a vast untapped resource in novel tools to modulate and perturb host cell processes, potentially via entirely novel targets and mechanisms of action. As *evolutionarily optimized perturbagens*, they might moreover be more effective in modulating cellular functions than more traditional perturbations such as knocking out or overexpressing human genes.

Several previous studies have characterized host protein interactions or perturbations by multiple pathogen effectors, but they have mostly covered a single pathogen species or relatively small collections of effectors.^24–34^ Thus, the discrepancy between the number of sequenced effector genes and those with known functions constitutes a major and critical knowledge gap for both fundamental discovery and potential therapeutics. To address this gap, we present a novel approach for investigating at an unprecedented scale how diverse effectors hijack host cell functions. Rather than focusing on a single pathogen or pathway, we developed scalable assays that enable the systematic functional profiling of thousands of effectors from a broad range of pathogens across a spectrum of host cellular pathways. This approach allowed us to uncover novel effectors that modulate key host proteins and provided general lessons for such approaches. Specifically, we uncovered novel functions for entirely uncharacterized ORFs, unexpected additional functions for known effectors, examples of close homologs with identical and with strongly divergent functions, as well as functions for recently evolved viral ORFs affecting a wide variety of pathways, including the NF-κB and p53 pathways, MHC-I antigen presentation, cGAS-STING signaling, and apoptosis. This work introduces a powerful framework for the annotation of host perturbations elicited by pathogen effectors and the discovery of novel protein function more generally.

## RESULTS

### The effector ORFeome collection 1.0

To systematically discover the function of thousands of eORFs, we assembled a large collection of effector ORFs. Currently, this collection, which we call the effector ORFeome (eORFeome), contains 3,835 putative effector ORFs (eORFs) from viruses, bacteria, and parasites (**Figure 1B, Figure S1** and **Table S1**). The collection includes virus-encoded ORFs from a broad phylogenetic range of viruses with representatives from all Baltimore classes, including dsDNA viruses (e.g. *Adenoviridae*, *Herpesviridae* and *Poxviridae*), ssDNA viruses (e.g. *Parvoviridae*, *Anelloviridae*), dsRNA viruses (e.g. *Reoviridae*, *Polyomaviridae*) and ssRNA viruses (e.g. *Coronaviridae*, *Orthomyxoviridae*, *Picornaviridae*). Our collection includes any virus-encoded ORF, including those encoding structural proteins. For polyproteins that are proteolytically processed into individual polypeptides, we included clones that correspond to the final cleaved proteins. For bacteria, the eORFeome comprises ORFs from curated lists of effectors secreted into host cells by type III, type IV or type VI secretion systems, such as those from *Legionella*, *Coxiella*, and *Chlamydia*.^35–37^ We also included eORFs from the human microbiome effector ORFeome,^38^ which encompasses effectors from commensal microbes from the human gut. Finally, we included 20 ORFs encoding predicted or known secreted effectors from the parasite *Toxoplasma gondii*.^39^ In total, the eORFeome contains 3,835 effectors from 25 viral, 12 bacterial and 1 parasitic family, representing a comprehensive collection of diverse pathogens and their niches.

The eORFeome exhibits a significant sequence diversity: the 3,835 eORFs cluster into 3,208 groups at 80% sequence identity and 2,118 groups at 10% identity (**Figure 1C**). We recently showed that even highly similar effectors can have distinct interaction profiles and hence likely different functions^38^. Therefore, for certain pathogens (e.g. HPV, HAdV, Enteroviruses), we included highly similar homologous ORFs from multiple strains (**Figure S1**). Notably, only 7% of the eORFs have direct experimental evidence for protein function and structure (i.e. UniProt annotation score of 5) compared to 72% of the human proteome (**Figure 1D**). The eORFs also are diverse in length, ranging from 22 to 2,225 amino acids (aa), with an average size of 345.5 aa (**Figure S2A**). Collectively, the eORFeome represents a functionally underexplored resource and a unique opportunity for large-scale characterization of pathogen effectors.

### A barcoded lentiviral library and platform for inducible eORF expression and high-throughput functional screening

We took advantage of the eORFeome to systematically identify pathogen effectors that hijack cellular functions using high-throughput selection-based screens. Inspired by screens based on human ORFeome overexpression,^22,23^ we constructed a lentiviral eORFeome library in an all-in-one vector that enables both doxycycline (dox)-inducible expression and detection of barcoded eORFs by next-generation sequencing (**Figure S2B**). Briefly, each eORF is followed by a unique DNA barcode that allows identification of the eORF independent of the eORF length and, moreover, increases the statistical power of screens due to an average of 200 unique barcodes per eORF (**Figure S2C** and **S2D**). Introducing this library into a starting cell population at low multiplicity of infection (MOI) ensures that most cells express only a single eORF. Controlled eORF expression by dox treatment coupled to phenotypic selection of cells by either reporter gene expression and FACS-based sorting or proliferation will enrich for cells that express eORFs with the respective function of interest. Next-generation sequencing of the barcodes allows the identification of enriched or depleted eORFs in selected cells compared to the starting population.

### A proof-of-principle screen for eORFs modulating the NF-κB pathway

The NF-κB pathway regulates genes controlling host immunity, cell proliferation and apoptosis, making it a common target for viruses, bacteria and parasites.^40–44^ Pathogens are known to inhibit NF-κB to suppress host immune response or, conversely, activate the pathway to avoid apoptosis (**Figure 2A**).^40^ Thus, the NF-κB pathway provides an ideal setting for a proof-of-principle screen of the eORFeome.

**Figure 2.**
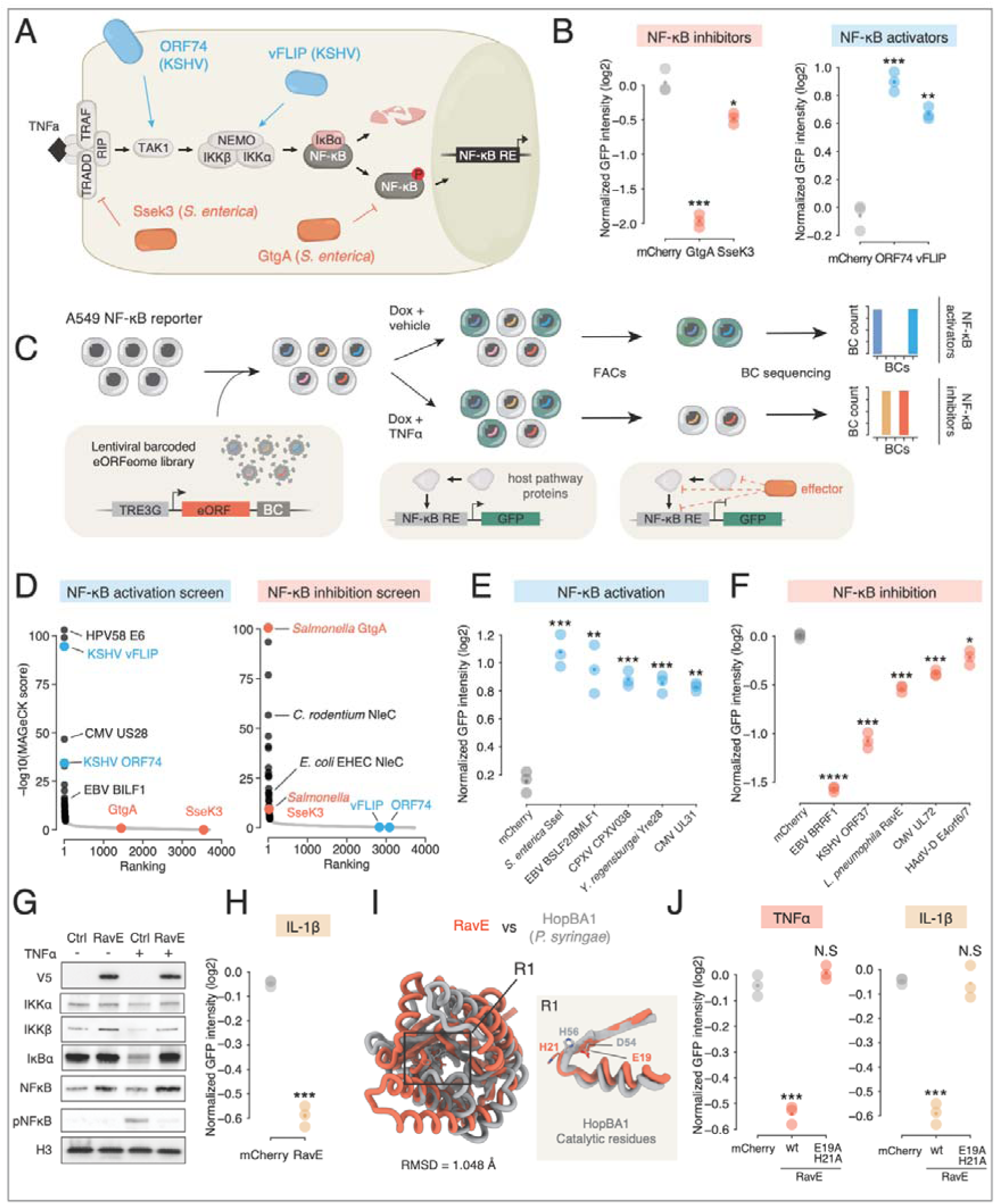
A proof-of-principle screen identifies known and novel NF-κB modulators. (A) Schematic of the NF-κB pathway. Points of intervention are shown for known inhibitory eORFs (S. enterica Ssek3 and GtgA) and activating eORFs (KSHV vFLIP and ORF74). (B) GFP intensity in an A549 NF-κB reporter cell line. Expression of the indicated ORFs was induced with dox. (Left) Cells expressing GtgA and SseK3 from Salmonella enterica, following treatment with 20 ng/mL TNFα for 16 hours to induce reporter activity. (Right) Cells expressing ORF74 and vFLIP from KSHV without TNFα treatment. The plots show the median GFP intensity, measured by flow cytometry and normalized to parallel cultures that were not treated with dox. The mCherry ORF serves as the negative control. Statistical significance was determined by comparing each ORF to the mCherry control using multiple paired t-tests. Dots indicate the mean of replicates. (C) Schematic of the screening platform for identifying eORFs that modulate NF-κB. The A549-NF-κB reporter cell line was used to monitor pathway activity. The reporter population was transduced at a low multiplicity of infection with a dox-inducible, barcoded lentiviral eORFeome library, ensuring single eORF expression per cell. For the activation screen (top), eORF expression was induced with dox, and GFP cells (indicative of pathway activation) were enriched by FACS. For the inhibition screen (bottom), eORF expression was induced with dox, followed by stimulation with TNFα (20 ng/mL) for 16 h, and cells with reduced GFP fluorescence were enriched by FACS. For both screens, eORF-associated barcodes from sorted and input populations were recovered from gDNA and quantified by deep sequencing. MAGeCK analysis was used to aggregate enrichment scores to systematically identify candidate eORFs with significant activating or inhibitory effects on NF-κB activity. (D) Ranking plot of enrichment scores for eORFs from the NF-κB activation (left) and inhibition (right) screens. Candidate hits (black dots) are distinguished from non-hits (grey) by a threshold of FDR < 0.05 and a fold change > 1.5. Positive controls for activation (vFLIP, ORF74; blue) and inhibition (Ssek3, GtgA; orange) are indicated, and other known regulators examples are labelled. (E) Validation of candidate eORFs from the NF-κB activation screen: SseI (*S. enterica*), BSLF2+BMLF1 (EBV), CPXV038 (Cowpox virus), Yre28/WP_006817680.1(*Y. regensburgei*) and UL31 (HCMV). The assay was performed under basal conditions as in (B, right). (F) Validation of candidate ORFs from the NF-κB inhibition screen: BRRF1 (EBV), ORF37 (KSHV), RavE (*L. pneumophila*), UL72 (HCMV), and E4orf6/7 (HAdV-D). The assay was performed with TNFα stimulation as in (B, left). (G) Western blot analysis of NF-κB pathway proteins in cells expressing the novel inhibitor RavE from L. pneumophila. Cells expressing V5-tagged RavE or an mCherry control were treated with or without TNFα (20 ng/mL) for 30 min. Lysates were immunoblotted with the indicated antibodies. An anti-V5 antibody was used to confirm RavE expression, and H3 served as a loading control. (H) Inhibitory activity of RavE assessed following pathway activation with IL-1β (1 ng/mL) for 16 h. (I) The structured domain of RavE (aa 1-234; orange) is shown aligned with the structure of the P. syringae effector HopBA1 (grey). A highlighted region, R1, indicates the alignment of catalytic residues from HopBA1 and RavE. The Root Mean Square Deviation (RMSD) indicates the degree of structural similarity between the two proteins. (J) Comparison of the inhibitory activity of wild-type RavE and a RavE (H21A,E19A) mutant following pathway activation with either TNFα (left) or IL-1β (right). ****p < 0.0001, ***p < 0.001, **p < 0.01, *p < 0.05; N.S, not significant. See also Figure S3 for representative examples of the single-cell GFP fluorescence distribution.

We engineered an A549 reporter cell line in which an NF-κB response element drives GFP expression (see Methods). We tested the ability of the reporter to detect NF-κB inhibition by pathogen effectors by expressing two known secreted NF-κB inhibitors from *Salmonella.* GtgA cleaves the NF-κB p65 subunit, whereas SseK3 inhibits the pathway by covalently modifying the TRADD receptor with N-acetylglucosamine.^43,45^ As expected, both eORFs strongly reduced GFP signal after stimulating the NF-κB pathway with TNFα (**Figure 2B** and **S3A**). Moreover, the reporter was induced in the absence of TNFα by two known viral NF-κB activators with distinct cellular targets and mechanisms: Kaposi’s sarcoma herpesvirus (KSHV) vFLIP that activates the IκB kinase (IKK) complex, and KSHV ORF74, a ligand-independent vGPCR that activates the TAK1 kinase (**Figure 2B** and **S3A**).^46,47,47,48^ These results validate the suitability of the reporter for NF-κB modulator screens.

To identify both inhibitors and activators, we transduced the reporter cells with the lentiviral eORFeome library and conducted two parallel screens (**Figure 2C**). In the inhibitor screen, we stimulated the pathway with TNFα and used FACS to isolate cells with low GFP levels. In the activator screen, we omitted TNFα and isolated cells with high GFP expression. Following isolation, we quantified the eORF-associated barcodes in each population by deep sequencing and used MAGeCK (see Methods) to compute enrichment scores.^49^

In total, the screens identified 29 activators and 42 inhibitors of NF-κB at a false discovery rate (FDR) of 0.05 and fold change > 1.5 (**Table S2** provides enrichment and FDR values for all eORFs), including the positive controls (**Figure 2D**). The screens recovered multiple known NF-κB modulators (**Figure 2D**). These included viral and bacterial effectors, such as NleC proteins from enterobacteria and 3C proteases from enteroviruses, which inhibit NF-κB by directly cleaving the p65 subunit.^50–52^ We identified three ligand-independent vGPCRs that mimic host chemokine receptors to activate NF-κB: Epstein-Barr virus EBV BILF1, cytomegalovirus (CMV) US28 and CMV UL138.^53–55^ Additionally, we found two glycoproteins implicated in NF-κB activation: HHV2 gD, a homolog of the NF-κB activator from HHV1,^56^ and EBV gH, which activates the pathway via interaction with TLR2.^57^ The activation screen also enriched for E6 proteins from multiple HPV types, which are known to induce NF-κB activity through an unknown mechanism.^58^

The screens also identified novel NF-κB regulators. To experimentally validate these candidates, we cloned a subset of novel hits and individually transduced them into the NF-κB reporter cell line. All five tested novel activator candidates strongly induced the NF-κB reporter in the absence of TNFα (**Figure 2E**). Furthermore, all five novel inhibitor candidates significantly suppressed NF-κB reporter activation, with four showing strong inhibition and one displaying weaker but still statistically significant inhibition compared to control (**Figure 2F**).

We selected the novel inhibitor RavE to elucidate the mechanisms by which it inhibits the NF-κB pathway activity. RavE (Lpg0195) is a Type IV Secretion System (T4SS) effector from *Legionella pneumophila* with an unknown domain architecture and function. RavE expression in A549 cells prevented the degradation of IκBα, the cytoplasmic inhibitor of NF-κB, and the subsequent phosphorylation of NF-κB (**Figure 2G**) after TNFα stimulation. Furthermore, RavE also inhibited IL-1β-mediated activation of NF-κB (**Figure 2H**). Since both the TNFα and IL-1β pathways converge at the IKK complex,^59^ these results suggest that RavE targets the step of IKK complex activation and IκBα degradation to block signaling rather than an upstream event.

Foldseek^60^ revealed RavE structural similarity with TIKI superfamily proteins^61^ and, in particular, with the *Pseudomonas syringae* effector HopBA1 that can activate host cell death in plants (**Figure 2I**).^62^ The TIKI superfamily comprises diverse esterases and proteases that share a putative active site.^61^ Structural alignment suggested that RavE residues E19 and H21, which are conserved among RavE homologs in other *Legionella* species (**Figure S3B**), might be catalytic residues. Indeed, mutating these residues to alanines completely abolished RavE’s ability to inhibit NF-κB activation by either TNFα or IL-1β (**Figure 2J and S3C,D**). Thus, RavE represents a novel Legionella effector that potently inhibits NF-κB pathway through highly conserved residues. More broadly, these results establish the eORFeome screens as a powerful screening and discovery platform to deliver assay-specific hits and uncover previously unknown functional host-pathogen interactions.

### eORFeome screens uncover known and novel effectors for four additional pathways

Building on the proof-of-principle NF-κB screens, we performed four additional screens representing pathways that are broadly relevant not only to infection but also to cancer, inflammation, and development: p53 signaling, STING signaling, apoptosis, and MHC-I antigen presentation. We selected these pathways because they play a central role both in normal cellular homeostasis and in host responses to infection. The p53 tumor suppressor regulates cell cycle progression, apoptosis, and DNA damage response and is a frequent target of viral oncogenes.^63^ The cGAS-STING pathway acts as the major innate immune sensor for cytoplasmic dsDNA. It is essential for detecting DNA viruses and intracellular bacteria, but it also plays an important role in tumor immunology and inflammation.^64^ Apoptosis is not only a conserved host defense mechanism against infection but also a key regulator of organismal development, tissue homeostasis and tumorigenesis.^65^ Finally, antigen presentation by MHC-I enables recognition of infected or transformed cells by cytotoxic T cells and thereby is a central target of immune evasion strategies employed by both pathogens and cancer cells.^66^

In each case, we employed well-established reporters or markers to identify pathway inhibitors. For p53, we used an A549 cell line with a p53 response element driving GFP expression (see Methods). After activating the pathway with the MDM2/p53 interaction inhibitor nutlin, we used FACS to sort low GFP cells. For STING, after activating the pathway with the STING ligand 2’,3’-cGAMP in U937 monocytes, we used an anti-IFIT1 antibody to sort cells that did not induce IFIT1 expression, a known cGAS-STING pathway target gene.^67,68^ We induced apoptosis in A549 cells with staurosporine followed by sequencing of eORF barcodes in the surviving population. Finally, for MHC-I surface display, we used a pan-MHC-I antibody and FACS to identify pathway inhibitors.

In each screen, we identified several known effectors, such as HPV31 E6 inhibiting p53, Vaccinia (VACV) poxin inhibiting STING signalling, EBV BHRF1 inhibiting apoptosis, or HHV7 U21 inhibiting MHC-I display (**Figure 3A, Table S2**). Moreover, the screens also uncovered hundreds of effectors with previously unrecognized activities, including both entirely uncharacterized proteins as well as known effectors with unexpected additional functions.

**Figure 3.**
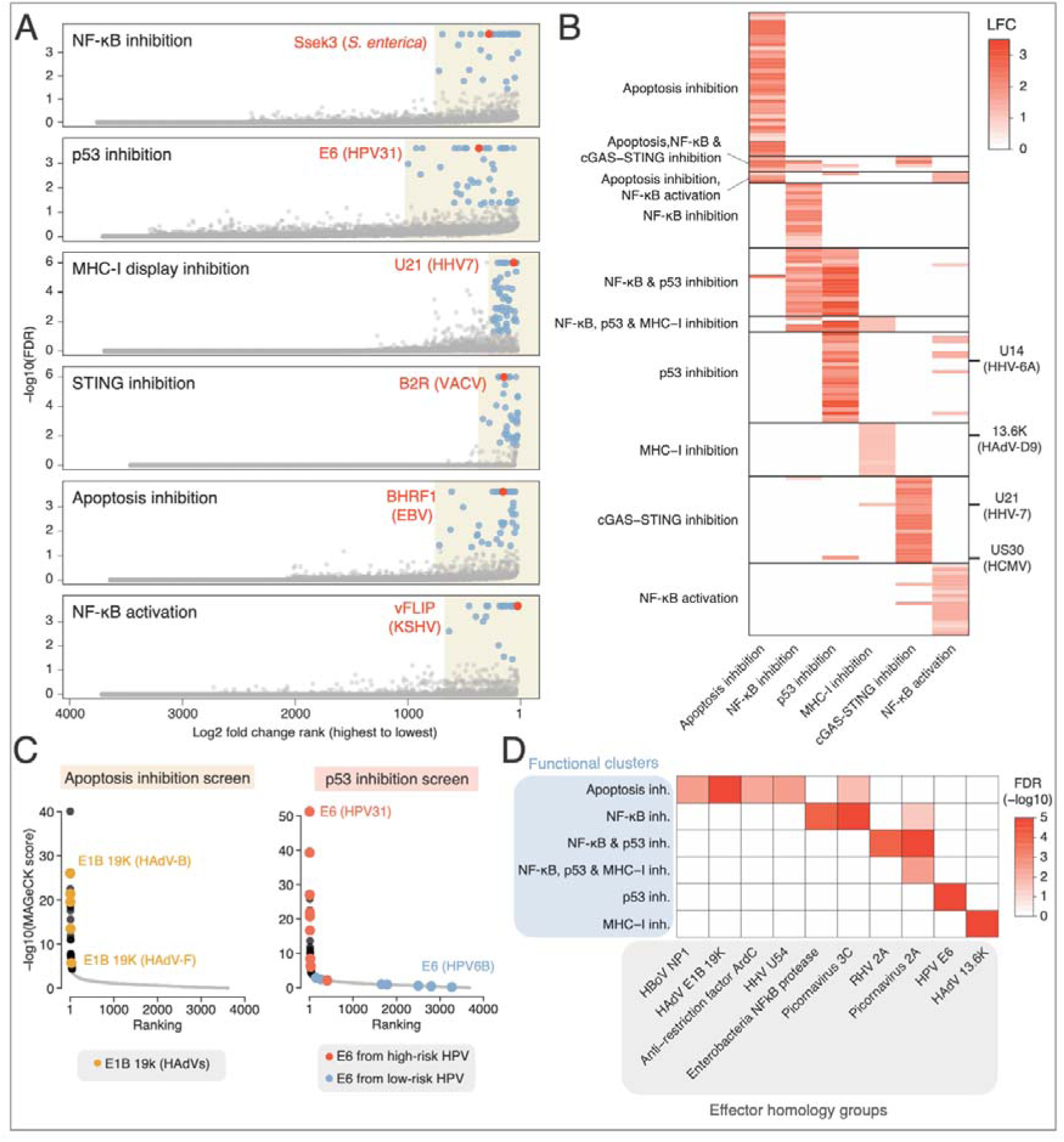
Parallel eORFeome screens uncover pathway-specific and pleiotropic effector functions. (A) Volcano plots with the results from six independent eORFeome screens. Each plot displays the enrichment (log2 fold-change; LFC) ranking on the x-axis (from highest/less enriched to lowest/more enriched) against the statistical significance (-log10 FDR) on the y-axis. Individual eORFs are represented as dots. Significant hits (FDR < 0.05 and FC > 1.5) are highlighted in blue, while a known pathway regulator is marked in orange for each screen as a positive control. (B) Self-organizing map (SOM) clustering of all significant hits (FDR < 0.05, LFC > 1) from the screens. The analysis groups eORFs into 10 distinct clusters based on their enrichment profiles (log-fold change values) across the different screens. (C) Enrichment ranking plot of eORFs from the apoptosis-inhibition screen (left) or p53-inhibition screen (right). Significant hits are marked in black. E1B 19K (HAdVs), E6 (high-risk HPVs) and E6 (low-risk HPVs) protein homologs are highlighted. (D) Enrichment of protein families (homology groups) within the functional clusters from (B). Homology groups were defined by clustering ORFs with a 30% sequence identity cutoff (MMseqs2^106^). The heatmap displays the FDR values for the enrichment of each homology group within each functional cluster. Only homology groups containing more than one ORF and with a significant enrichment (FDR < 0.05) in at least one functional cluster are shown.

### Parallel eORFeome screens uncover pathway-specific and pleiotropic effectors as well as shared and distinct functions for closely related eORF homologs

Our screens revealed 164 distinct effector hits across the six screens. Although the screens targeted distinct pathways, 24% (40/164) of the hits scored in multiple screens, suggesting that a large fraction of pathogenic effectors target multiple processes. To discriminate pathway-specific from more pleiotropic effects, we used self-organizing map clustering to group the hits found in any of the screens into ten groups with distinct patterns of high to low scores in one or several screens (**Figure 3B, Table S3**). Most clusters were specific to a single pathway. Suggesting conserved functions for pathogen effectors, the functional clusters uncovered multiple homologous proteins among screen hits. For example, the apoptosis inhibition screen identified multiple Bcl2-like proteins, including five homologs of adenoviral E1B proteins known to suppress apoptosis (**Figure 3C**).^69^ Similarly, among the top hits of the p53 screen were several E6 proteins from papillomaviruses that are associated with a high risk of cervical cancer or head and neck cancer. In contrast, none of the E6 proteins from low-risk HPV strains were identified as screen hits (**Figure 3C**), consistent with the well-established oncogenic role of E6 from high-risk strains in degrading host p53.^70,71^ Thus, the screens identified both conserved and strain-specific phenotypes for effectors, highlighting the power of unbiased, sensitive screens in functional discovery.

We identified several protein families (defined as homology groups of eORFs sharing at least 30% sequence identity) that were enriched in specific clusters (adjusted p-value < 0.05, **Figure 3D)**. Some protein families were already linked to their respective cellular pathway, including the NF-κB-degrading zinc proteases from enterobacteria,^43,72,73^ E6 proteins from HPVs^70,71^ and E1B proteins from HAdV.^69^ However, involvement of other protein families – including HHV U54 tegument proteins in apoptosis, and adenoviral i-leader/13.6K proteins in MHC-I display – has not been reported and could not have been predicted e.g. by sequence similarity. Thus, parallel eORFeome screens can functionally characterize hundreds of pathogen effector proteins, uncovering not only novel activities for individual effectors but also previously unrecognized functions for entire protein families as well as additional functions for known effectors. Next, we set out to validate and characterize novel hits corresponding to each of these general lessons, selected from three different cellular pathways.

### Discovery of a novel and unexpected function: HHV6A U14 is a potent p53 antagonist

One of the most prominent hits in our p53 screen was the U14 protein from human herpesvirus 6A (HHV6A) **(Figure 4A**). A prior study reported that U14 interacts with p53 but concluded that it does not inhibit p53 activity.^74^ We therefore investigated U14 in more detail to understand its mechanism of action and to assess if it is a true p53 antagonist.

**Figure 4.**
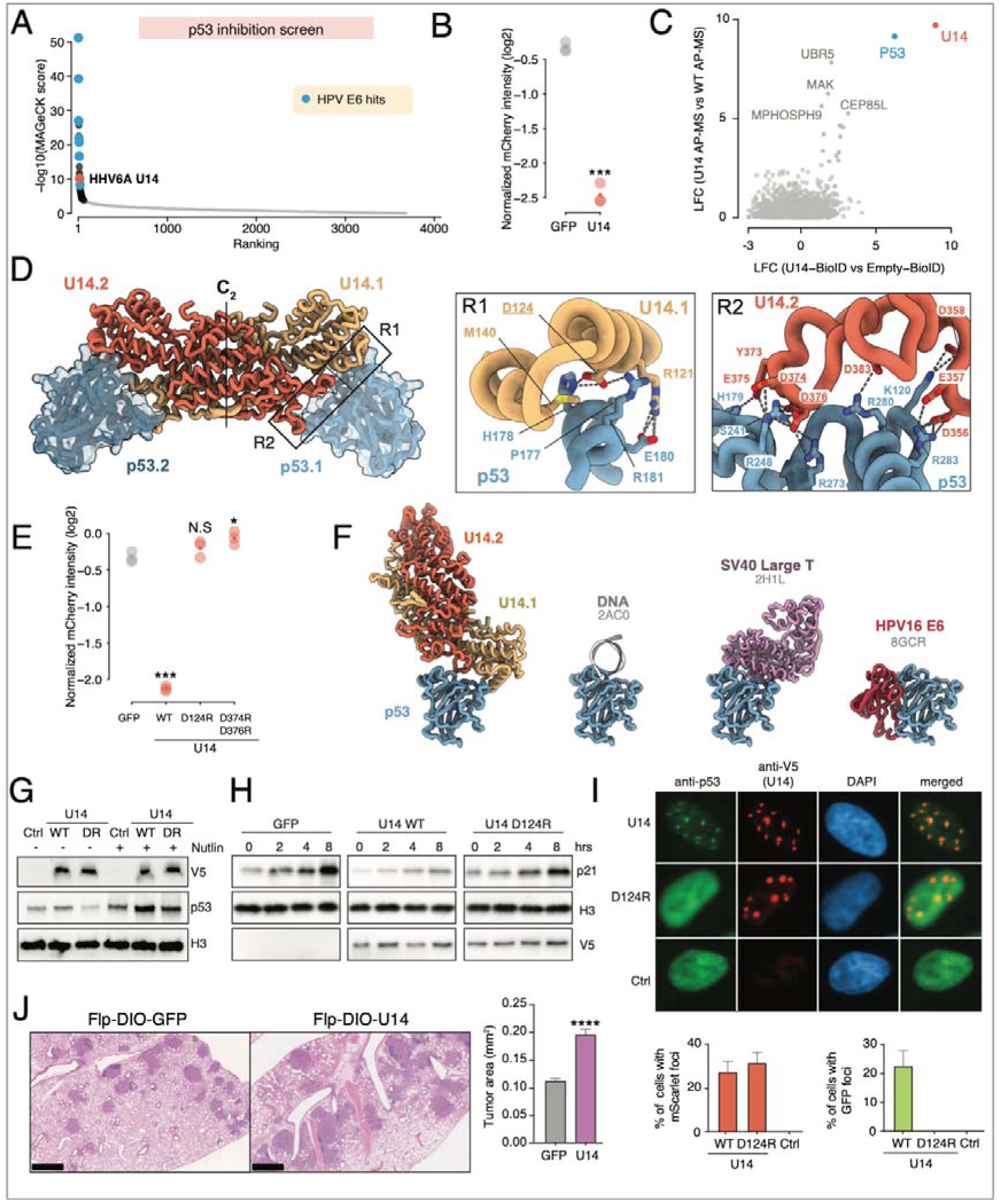
U14 from HHV6A is a potent p53 antagonist. (A) Enrichment ranking plot of eORFs from the p53-inhibition screen using a p53-mCherry reporter in A549 cells. Significant hits (FDR < 0.05 and FC > 1.5) are highlighted in black. HPV E6 proteins that scored as hits are highlighted in blue; U14 from HHV[6A is highlighted in orange. All other eORFs are shown in grey. (B) mCherry intensity in an A549 p53 reporter cell line. Expression of the novel candidate U14 (HHV-6A) was induced with dox, following treatment with nutlin (2.5 µM) for 16 hours to induce reporter activity. The plots show the median mCherry intensity, measured by flow cytometry and normalized to parallel cultures that were not treated with dox. The GFP ORF serves as the negative control. Statistical significance was determined by comparing each ORF to the GFP control using multiple paired t-tests. Dots indicate the mean of replicates. (C) Scatter plot of protein enrichment scores from BioID (x-axis) and IP-MS (y-axis) for U14 relative to an empty control. (D) AlphaFold 3 model of the U14-p53 complex. The U14 dimer is shown in orange and yellow, bound to two p53 proteins in blue and light blue. Black rectangles (R1 and R2) mark the interaction regions. The right panels show enlarged views of the interaction regions R1 and R2 from the structural model. Key amino acid residues are shown and labelled. (E) Comparison of the inhibitory activity of wild-type U14, U14 D124R mutant or U14 D374R,D376R mutant following pathway activation with nutlin (2.5 µM) for 16 hours. (F) Structural models of p53 in complex with various binding partners. Shown are PDB structures of p53 bound to DNA,^123^ SV40 LTAg^75^ and HPV E6,^124^ alongside an AlphaFold 3 model of the predicted interaction between U14 dimers and p53. (G) Western blot analysis of cells expressing GFP, V5-tagged U14, or V5-tagged U14 D124R mutant, treated with or without 2.5 µM nutlin for 1 hour. Blots were probed with antibodies against V5, p53, and Histone H3 as a loading control. (H) Western blot analysis of cells expressing GFP, V5-U14, or V5-U14 D124R, treated with nutlin for the indicated times (0-8 hours). Blots were probed for p21, V5, and H3 loading control. (I) Immunofluorescence of cells expressing an empty control, V5-tagged U14, or V5-tagged U14 D124R. Cells were stained with antibodies against p53 and V5, with DAPI for nuclear visualization. Bottom: Quantification of the percentage of cells with p53 foci (left) and U14 foci (right). Data are shown as mean ± SD from three independent experiments. (J) In vivo evaluation using a Rosa-CreERT2; FSF-KrasG12D lung adenocarcinoma mouse model. Lentivirus co-expressing U14 or GFP control was delivered via inhalation, followed by tamoxifen induction. Lungs were harvested 9 weeks later, H&E stained, and tumour area quantified. FACS data in all panels is shown as mean ± SD from three independent experiments. Statistical significance is calculated relative to the mCherry (negative control) + dox condition for all panels, unless otherwise specified by lines indicating alternative comparisons, and is denoted as follows: *padj < 0.05, **padj < 0.01, ***padj < 0.001, **** padj < 0.0001. See also Figure S4 for representative examples of thesingle-cell mCherry fluorescence distribution.

We expressed U14 in a p53 reporter cell line and observed a marked suppression of p53 reporter activation, validating U14’s role as a potent p53 antagonist (**Figure 4B**). Additionally, we validated the physical interaction between p53 and V5-tagged U14 with both affinity-purification coupled to mass spectrometry (AP-MS) and proximity-dependent biotinylation (BioID) (**Figure 4C and Table S4**). AlphaFold2 predicted that U14 forms a dimer with two principal regions of p53 contact (**Figure 4D** and **S4A**): Region 1 (R1) comprises residues D124 and R121 of one U14 protomer, predicted to interact with the core domain of p53 residues 177–181. Region 2 (R2) includes residues 356–376 of the second U14 protomer that are positioned to interact with the DNA-binding domain of p53 (aa 241–283). Thus, it is likely that both regions interfere with the binding of p53 to its cognate DNA elements. Consistent with these predictions, point mutations in R1 (D124R) or R2 (D374R, D376R) abolished U14’s ability to inhibit p53 activity (**Figure 4E** and **S4B,C**), thus supporting the structural model and highlighting the critical role of each of the two regions.

The U14-p53 interaction surface is also targeted by the Large T antigen (LTAg) from SV40,^75^ a well-studied p53 antagonist (**Figure 4F**) that is unrelated to U14. LTAg stabilizes and inhibits p53 DNA binding,^76^ in contrast to HPV E6 that degrades p53.^70,71^ In line with the shared interaction surface, U14 did not reduce p53 protein levels, but rather stabilized p53 (**Figure 4G**). Furthermore, U14, but not the D124R mutant, impaired p21 induction following nutlin treatment in A549 cells (**Figure 4H**). Strikingly, U14 expression also led to p53 sequestration into distinct nuclear foci that overlap with U14 localization (**Figure 4I**). This relocalization depends on direct U14-p53 interaction, as the U14-D124R mutant, while still forming nuclear foci, could not recruit p53. Together, these findings suggest a functional evolutionary convergence between dissimilar effectors of two unrelated viruses, both capable of sequestering and stabilizing p53 while suppressing its trans-activation capacity.

Next, we hypothesized that U14 would interfere with p53-mediated tumor suppression and cooperate with the Kras^G12D^ oncogene to drive tumor formation.^77^ To test U14’s biological impact *in vivo*, we engineered lentiviral vectors that contain the FLP recombinase and a double-floxed inverted orientation (DIO) cassette of U14 or eGFP (LV-Flp-DIO-U14 or LV-Flp-DIO-eGFP), which we delivered through intranasal inhalations into the lungs of Frt-Stop-Frt (FSF)-Kras^G12D^; Rosa26-CreERT2 mice^78,79^, where a premature stop codon prevents Kras^G12D^ expression in the absence of recombination. FLP recombinase clonally activates the FSF-Kras^G12D^ allele within the lung of inhaled mice, while administration of tamoxifen allows temporal control of the expression of U14 or eGFP via CreERT2. In alignment with our hypothesis, U14-expressing mice developed nearly double the tumor area compared to the eGFP controls (**Figure 4J**). This result provides strong evidence that U14 functions as a potent p53 antagonist in vivo, accelerating tumorigenesis by dismantling the cell’s primary defense against oncogenic transformation.

### Discovery of a new function for a known viral effector: HHV7 MHC-I inhibitor U21 is also a potent inhibitor of STING signaling

Similar to the p53 screen, our screen for STING pathway inhibitors identified multiple known modulators of the pathway. One of the top hits was VACV poxin, which was recently described as a nuclease that degrades cGAMP (**Figure 5A**).^80^ Other known STING pathway inhibitors included CMV UL42, which inhibits STING translocation,^81^ and the Rabies virus P phosphoprotein that inhibits TBK1 and IRF3 function downstream of STING.^82,83^ These results suggested that other hits would also be physiologically relevant modulators of the pathway. In particular, among the hits were two herpesvirus proteins, CMV US30 and HHV7 U21, with no previous connection to cGAS-STING signaling.

**Figure 5.**
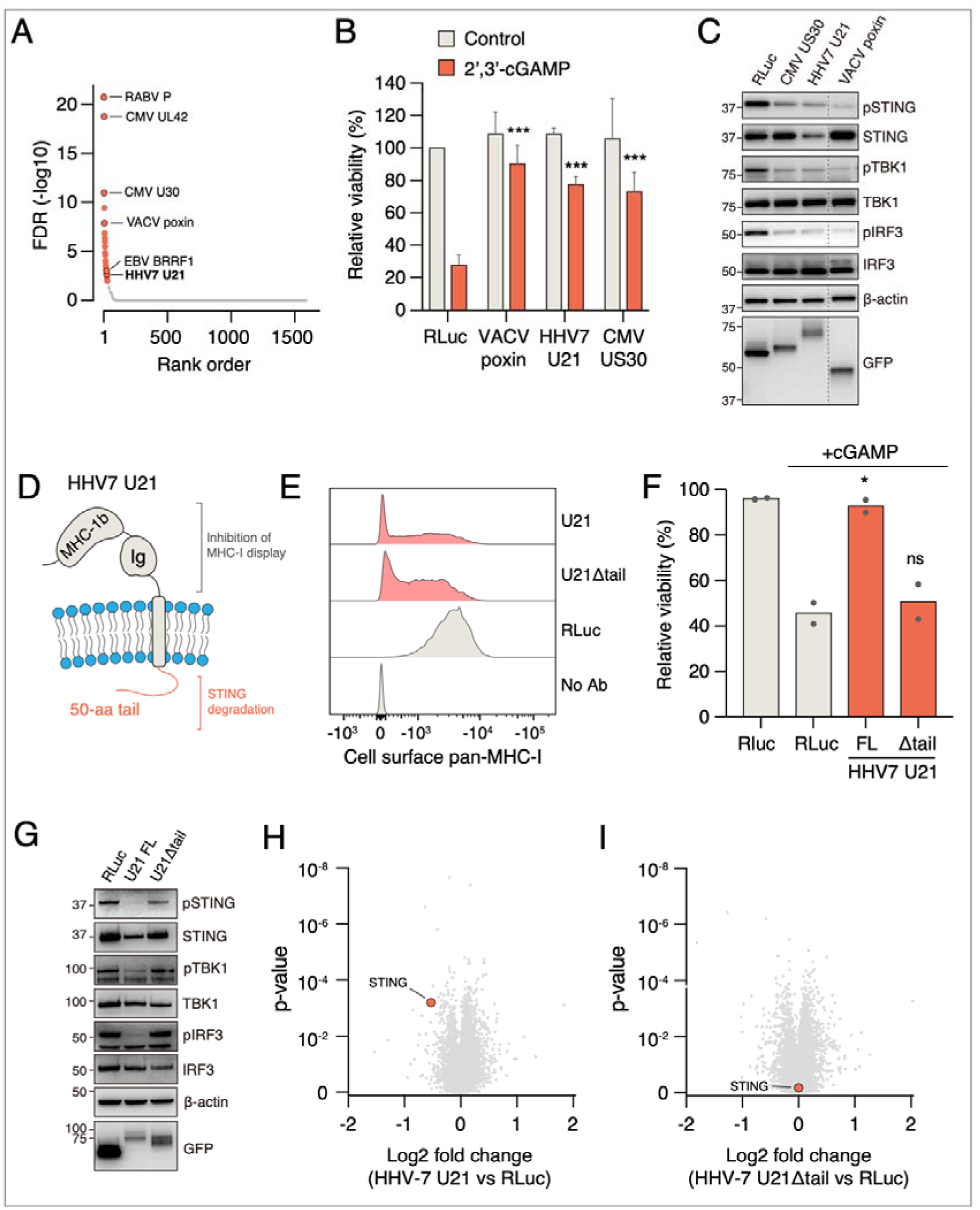
HHV7 U21 antagonizes cGAS-STING signaling by inducing STING degradation. (A) Enrichment ranking plot of eORFs from the STING inhibition screen in U937 cells. Significant hits are highlighted in red. Notable known inhibitors of STING signaling are labeled, with U21 highlighted as a novel STING inhibitor. All other eORFs are shown in grey. (B) U937 cells infected with indicated lentiviral dox-inducible GFP-tagged constructs, treated with dox and 40 µg/ml 2’,3’-cGAMP or vehicle and assessed for viability with CellTiter-Glo. (C) Indicated GFP-tagged proteins were expressed in cGAMP-treated U937 cells followed by western blotting for STING pathway components. (D) Schematic of HHV7 U21 protein. (E) U937 cells expressing full-length U21 (FL), U21 lacking the C-terminal tail (Δtail, aa 1-380), or Renilla luciferase were stained for cell surface MHC-I and analyzed by flow cytometry. (F) Full-length U21 and U21Δtail were assessed for suppression of cGAMP-induced cell death as in (B). (G) Full-length U21 and U21Δtail were assessed for inhibition of STING pathway by western blotting as in (C). (H-I) Full-length HHV7 U21 (H) or U21Δtail (I) were expressed in U937 cells, which were subjected to whole cell proteomics.

Activation of STING signaling with STING agonists in U937 cells leads to cell death that is dependent on an intact STING pathway.^84,85^ We therefore assessed the ability of dox-inducible, GFP-tagged VACV poxin, CMV US30, and HHV7 U21 to suppress cGAMP-induced cell death. Consistent with the screen, all proteins potently suppressed cGAMP-induced apoptotic cell death when expressed individually (**Figure 5B** and **S5A**). However, in contrast to poxin, which could suppress cell death induced by cGAMP but not by the non-cyclic dinucleotide STING agonist diABZI,^86^ the other two viral effectors counteracted the effect of both ligands (**Figure S5B**). Thus, the novel effectors function through a different mechanism than Vaccinia poxin.

All three effectors also inhibited STING phosphorylation as well as phosphorylation of downstream signaling components TBK1 and IRF3 after cGAMP treatment (**Figure 5C**). Interestingly, however, US30 and U21 had different effects: US30 inhibited STING phosphorylation without affecting its levels, whereas U21 expression led to lower STING protein levels. U21 is a single-pass transmembrane domain protein with an MHC-Ib-like and Ig-like lumenal domains and a 50 amino acid unstructured cytoplasmic tail (**Figure 5D**).^87^ U21 was originally identified as an immunoevasin that routes MHC-I molecules to lysosomes to evade recognition by cytotoxic T cells.^88,89^ The lumenal domain is required for MHC-I redirection to lysosomes but the cytoplasmic tail is dispensable for this activity,^90^ which we also confirmed (**Figure 5E**). However, in contrast to this, we observed that the cytoplasmic tail was indispensable for STING pathway inhibition, indicating that these activities are genetically separable. Deleting the tail abolished the ability of U21 to suppress cGAMP-induced cell death (**Figure 5F and S4C**), downregulate STING levels, and inhibit cGAMP-induced STING, TBK1, and IRF3 phosphorylation (**Figure 5G**). There was no difference in STING mRNA levels, ruling out a transcriptional downregulation (**Figure S5D**). Moreover, U21 could downregulate STING levels independent of cGAMP treatment, indicating that it modulates basal STING levels (**Figure S5E**).

To investigate if U21 induces protein degradation in a more general manner, we conducted a whole-cell proteomics experiment in U937 cells expressing full-length GFP-tagged U21, HHV7 U21 lacking the cytoplasmic tail (U21Δtail), or Renilla luciferase as a control. After inducing the expression for 24 hours with dox, HHV7 U21 significantly upregulated only 5 proteins and downregulated 5 proteins, including STING (**Figure 5H** and **Table S5**). In contrast, U21Δtail did not have an effect on STING levels (**Figure 5I**). Thus, U21 has a limited but very specific effect on the global proteome and it uniquely downregulates STING via its C-terminal tail. Furthermore, it represents a dual-function effector that can inhibit both adaptive and innate immune responses by targeting MHC-I molecules and STING, respectively, with topologically distinct domains.

### Functional annotation of a recently evolved non-canonical eORF: Adenoviral i-leader/13.6K proteins inhibit surface MHC-I expression via TAP transporter inhibition

Our MHC-I surface display screen identified multiple known regulators of MHC-I display. These included HHV7 U21 characterized above, CMV US11 and US2 proteins that dislocate MHC-I molecules from the ER to the cytosol for proteasomal degradation,^91,92^ and KSHV MIR2 E3 ligase that ubiquitinates and degrades MHC-I (**Figure 6A**).^93^ However, the most prominent novel hits were 13.6K proteins from four distinct human adenovirus serotypes HAdV-A12, HAdV-B21, HAdV-D9, and HAdV-F41 (**Figure 6A**).

**Figure 6.**
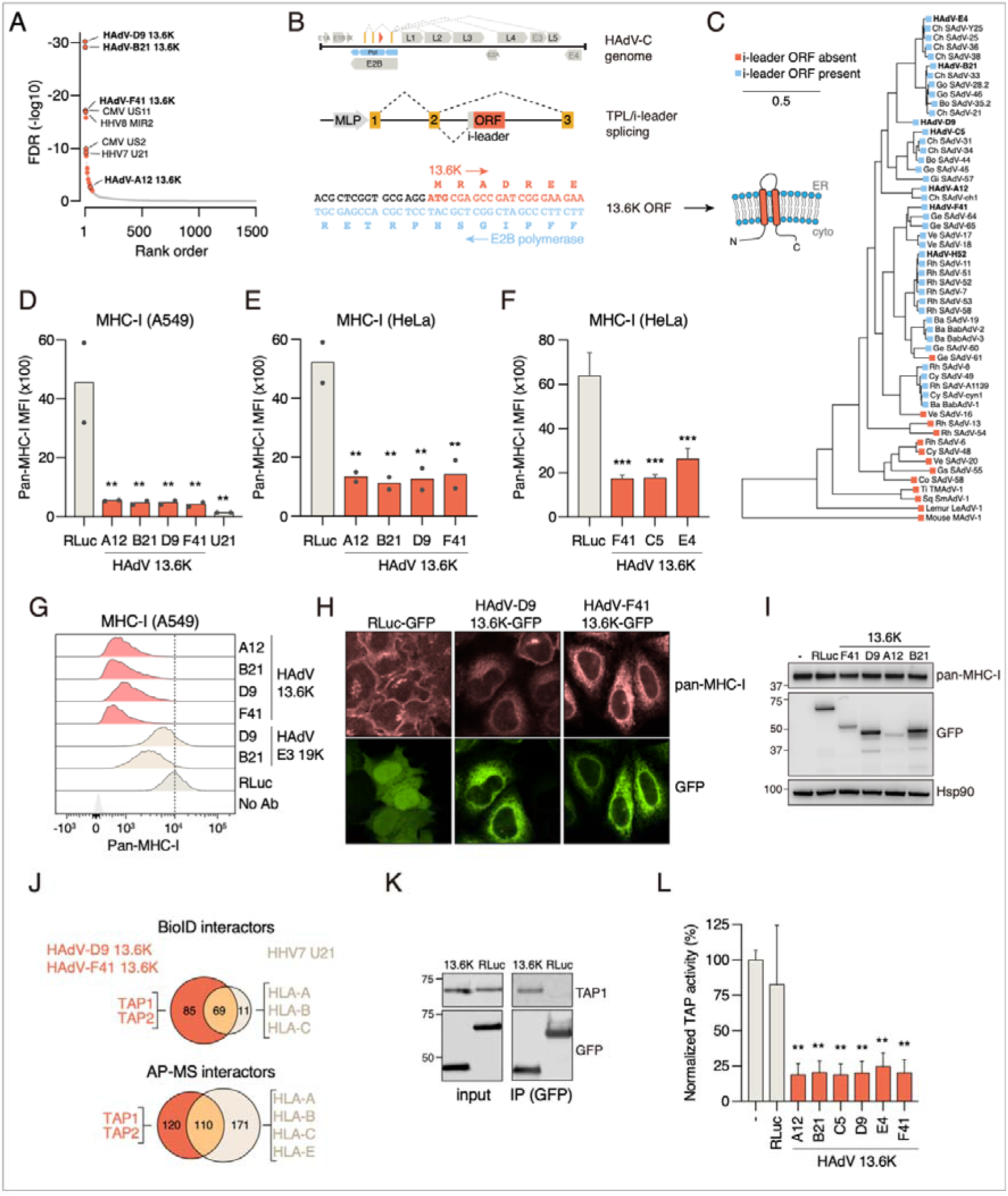
Adenoviral 13.6K/i-leader protein is a recently evolved inhibitor of the TAP transporter. (A) Enrichment ranking plot of eORFs from the MHC-I display screen in A549 cells. Significant hits are highlighted in red. Notable known inhibitors of STING signaling are labeled, with 13.6K clones present in the library highlighted as novel MHC-I regulators. All other eORFs are shown in grey. (B) 13.6K is encoded by the i-leader ORF located between the second and third adenoviral tripartite leader exons, on the opposite strand of the E2B polymerase. (C) Phylogenetic analysis of i-leader ORF presence in primate adenoviruses. Human adenoviruses are represented by a single strain from each clade whereas primate adenoviruses include representatives of 98% sequence identity clusters. Ch, chimpanzee; Bo, bonobo; Go, gorilla; Gi, gibbon; Ge, gelada baboon; Ve, vervet monkey; Rh, rhesus macaque; Ba, baboon; Cy, cynomolgus macaque; Gs, Golden snub-nosed monkey; Co, Colobus monkey; Ti, titi monkey; Sq, squirrel monkey. (D) A549 cells stably expressing dox-inducible GFP-tagged 13.6K proteins or HHV7 U21 were treated with dox for 24 hours followed by flow cytometry analysis of cell surface MHC-I levels. (E) The same 13.6K-GFP constructs were analyzed in HeLa cells as in (D). (F) 13.6K-GFP proteins from HAdV-F41 and HAdV-C5 were similarly analyzed in HeLa cells. (G) Comparison of MHC-I surface levels in A549 cells expressing 13.6K-GFP or the previously known immunoevasin E3 19K-GFP from HAdV-D9 or HAdV-B21. (H) HeLa cells expressing HAdV-D9 or HAdV-F41 13.6K-GFP or Renilla luciferase fused to GFP were stained for MHC-I. The GFP channel shows localization of the GFP-tagged constructs. (I) Western blot analysis of total MHC-I levels in HeLa cells expressing indicated 13.6K-GFP constructs. (J) HAdV-D9 or HAdV-F41 were tagged with miniTurbo-3xFLAG in HeLa cells and analyzed for physical and proximity interactors with BioID and AP-MS, respectively. HHV7 U21 served as a control for an immunoevasin that inhibits surface trafficking of MHC-I molecules. Venn diagrams show the overlap of 13.6K and U21 interactors. (K) HAdV-D9 13.6K-GFP or Renilla luciferase-GFP expressed in HeLa cells was immunoprecipitated with an anti-GFP antibody followed by western blotting for endogenous TAP1 or GFP. (L) GFP-tagged Renilla luciferase or 13.6K from indicated adenoviral serotypes were expressed in HeLa cells and semi-permeabilized followed by a translocation assay of fluorescently labeled TAP substrate peptide into the ER in the presence of ATP and Mg^2+^. Transported peptide was detected by flow cytometry.

The 13.6K protein (also known as i-leader protein) is encoded on the opposite strand of the essential E2B polymerase by an open reading frame located between the second and third exons of the tripartite leader (TPL).^94^ The TPL is a constant 5’ leader sequence for all late adenoviral transcripts (**Figure 6B**), and some leader splicing events include the intervening leader (i-leader) with the open reading frame encoding 13.6K (**Figure 6B**).^95^ 13.6K is predicted to have two transmembrane domains with both termini located on the cytoplasmic side (**Figure 6B**) but its function is unknown. Interestingly, the i-leader ORF is restricted to Old World monkey and hominid adenoviruses, as indicated by the absence of a start codon or the presence of in-frame stop codons in the i-leader ORF in other lineages (**Figure 6C**). Thus, 13.6K represents a recently evolved viral protein with an unknown function.

We validated that 13.6K proteins from the four human adenovirus strains identified in the screen downregulated MHC-I display in A549 and HeLa cells when expressed individually fused to C-terminal GFP (**Figure 6D** and **6E**). We also cloned the ORF from two additional serotypes (HAdV-C5, HAdV-E4) that were not included in the original screen. Both HAdV-C5 and HAdV-E4 13.6K robustly downregulated surface MHC-I, indicating that this is a conserved function of the protein (**Figure 6F**). 13.6K also inhibited surface display of β2 microglobulin, a constant subunit of all MHC class I molecules (**Figure S6A** and **S6B**). However, HAdV-F41 13.6K expression had no effect on the surface levels of two other plasma membrane proteins EGFR and CD47, suggesting that the effect is specific to MHC-I rather than 13.6K generally interfering with trafficking to the plasma membrane (**Figure S6C**). Moreover, 13.6K proteins were significantly more effective in MHC-I downregulation than the previously characterized adenoviral immunoevasin E3 19K from HAdV-B21 or HAdV-D9 (**Figure 6G**).^96^

13.6K could reduce MHC-I surface levels by interfering with membrane trafficking or by inducing MHC-I degradation. To differentiate between these options, we stained 13.6K-GFP or RLuc-GFP expressing HeLa cells with a pan-MHC-I antibody. In control cells expressing RLuc-GFP, MHC-I was localized to the plasma membrane as expected (**Figure 6H**). In contrast, in 13.6K expressing cells, there was no plasma membrane staining of MHC-I. Instead, we detected MHC-I in the endoplasmic reticulum together with 13.6K-GFP (**Figure 6H**). Moreover, 13.6K proteins did not affect total MHC-I levels as assessed by western blotting (**Figure 6I**), indicating that they are ER-associated membrane proteins that inhibit MHC-I membrane trafficking rather than stability (**Figure 6I**).

To understand how 13.6K disrupts antigen presentation, we characterized the interactomes of HAdV-D9 and HAdV-F41 13.6K in HeLa cells with AP-MS and BioID. We tagged them at the C termini with UltraID-3xFLAG and used HHV7 U21 as a positive control for an effector that downregulates MHC-I, and EGFP as a negative control. As previously reported, HHV7 U21 associated with HLA-A, HLA-B and HLA-C by both BioID and AP-MS, whereas 13.6K proteins did not significantly associate with these proteins (**Figure 6J** and **Table S5**). In contrast, HAdV-D9 and HAdV-F41 13.6K associated with the heterodimeric TAP peptide transporter complex subunits TAP1 and TAP2 in both BioID and AP-MS (**Figure 6J**). Endogenous TAP1 also co-immunoprecipitated with GFP-tagged HAdV-D9 13.6K, validating the interaction (**Figure 6K**). The TAP complex transports cytosolic peptides to the ER for loading them onto MHC-I molecules as part of the peptide loading complex (PLC). Many viral effector proteins are known to target TAP to inhibit MHC-I loading and trafficking to the membrane,^97^ making the TAP complex a strong candidate as the cellular target of 13.6K proteins.

To test if 13.6K proteins could directly inhibit TAP peptide transporter activity, we expressed EGFP-tagged 13.6K proteins from four serotypes (A12, B21, D9, and F41) in HeLa cells by lentiviral infection and assessed TAP activity with a fluorescent peptide transporter assay.^98^ We permeabilized cells expressing GFP-tagged 13.6K or Renilla luciferase with a mild detergent and incubated them with a fluorescently labeled TAP substrate peptide RRYQNSTC(AF647)L in the presence of ATP.^99^ In these conditions, TAP can transport the substrate peptide into the ER, which can be detected by flow cytometry. All 13.6K proteins significantly inhibited TAP activity (**Figure 6L**), establishing that they represent a novel family of viral TAP inhibitors.

## DISCUSSION

### The eORFeome as a discovery platform

The eORFeome constitutes a versatile discovery platform that bridges pathogen genomics and functional biology. By combining a comprehensive, sequence-diverse library of almost 4,000 viral, bacterial, and parasitic effector ORFs with barcoded, inducible expression and multiplexed phenotypic screening, this platform enables systematic mapping of effector activities across diverse host pathways. Unlike CRISPR-based perturbations (CRISPR-KO, CRISPRi, and CRISPRa), which rely on the one-dimensional modulation of endogenous gene activity within a single species, the eORFeome assays the functional potential of thousands of exogenous proteins optimized by evolution to rewire host processes in diverse, potentially multidimensional ways. This approach captures mechanisms inaccessible to genetic perturbation of host genes alone and complements CRISPR-based strategies by identifying exogenous regulators that mimic, oppose, or reveal latent host regulatory principles. Thus, eORFs exploit, as *evolved perturbagens,* a wide variety of cellular target proteins and protein-protein interfaces critical for host cell function and central to the modulation of host cell regulatory networks. The modular design of the eORFeome screening platform, encompassing pathway-specific reporters, survival-based selections, and scalable barcoded readouts, makes it adaptable to virtually any cellular process that can be measured at population scale.

### General lessons from diverse eORFeome screens

Functionally, the eORFeome screens illuminate general principles of pathogen-host interaction and, more broadly, of protein evolution and function. First, they establish at a systematic scale that entirely unannotated proteins, often with no structural or sequence similarity to known proteins, can execute highly specific and potent cellular functions. Second, many effectors are multifunctional, influencing distinct pathways through separable domains or interfaces, exemplified by herpesviral U21, which employs distinct regions to inhibit both adaptive immunity via suppressing MHC-I antigen presentation and innate immunity by inducing STING degradation. Third, closely related homologs can diverge sharply in activity, such as the high-risk versus low-risk HPV E6 proteins with distinct abilities to inhibit p53. Yet despite this divergent, rapidly evolving sequence space, diverse effectors recurrently converge on a limited number of host pathways and proteins, highlighting evolutionary “hotspots” of host vulnerability and suggesting that the host proteome might impose a finite set of exploitable interfaces.

### Specific findings from diverse eORFeome screens

Among the specific discoveries, our work identifies (i) U14 from HHV6A as a p53 antagonist that sequesters p53 without degrading it, revealing a mode of inactivation analogous to the unrelated SV40 large T antigen; (ii) the herpesviral U21 as a dual-function effector that blocks both antigen presentation and innate DNA sensing, via separable structural modules; and (iii) adenoviral 13.6K proteins as a novel family of TAP inhibitors that suppress MHC-I surface expression. The newly uncovered role of 13.6K proteins is a particularly illuminating example of the function and evolution of pathogenic effectors. 13.6K protein is expressed at high levels during infection,^100^ but no function had been identified. The ability of adenoviruses to evade the adaptive immune system has so far been attributed to the E3 19K immunoevasin that has been shown to interact with and inhibit the surface trafficking of MHC-I molecules. However, human adenovirus A, F, and G species lack E3 19K, leading to the suggestion that these species might evade host immunity by other means and targets.^101^ In contrast, 13.6K is present in all HAdV clades, is more abundant than E3 19K during infection,^102^ and is more potent in downregulating MHC-I surface presentation (**Figure 6G**), suggesting that it might be a more critical immunoevasin for adenoviral fitness. Strikingly, the 13.6K protein emerged during primate adenovirus evolution by a combination of a novel alternative splicing event in the tripartite leader, yielding the i-leader exon, and the acquisition of an ORF in the opposite strand but in the same reading frame as the highly essential E2B polymerase. This remarkable evolutionary path highlights how pathogens can evolve novel functional proteins *de novo* even within exceptionally highly constrained sequences. More generally, our findings not only define new effector classes and mechanisms but also extend the conceptual framework of viral evolution, showing how new ORFs can emerge through splicing innovation and quickly acquire immunomodulatory activity.

### Outlook and vision: a novel way of discovery

Our work defines a new paradigm for protein function discovery, one that treats pathogen effectors from across kingdoms as an evolutionary library of host modulators and functional building blocks, i.e. *evolved perturbagens*. By studying individual eORFs outside their pathogenic contexts, the approach decouples effector biology from disease models, enabling systematic discovery of molecular functions in a controlled and safe setting across evolutionarily unrelated pathogens. This cross-pathogen perspective not only accelerates annotation of previously uncharacterized proteins but also generates foundational data for using eORFs as molecular tools and for training emerging sequence-to-function models. Moreover, it provides an inspiration for engineering minimal domains or synthetic proteins that modulate host pathways with precision for next-generation therapeutics, pathway engineering, and beyond.

In the longer term, coupling large-scale functional genomics with eORF collections that span highly divergent, unannotated protein sequences will enable direct mapping from sequence to function and sequence-to-function protein models. Such data-driven modelling will establish a route toward predictive and design-based biology, where the landscape of natural effector mechanisms serves as a training ground for understanding, predicting, and ultimately engineering cellular control.

## ACKNOWLEDGEMENTS

We thank Tanisha Teelucksingh and Mandy Lam from the Taipale lab and Kevin Sabath, Kavya Shetty, Michaela Pagani, and Katharina Bergauer from the Stark lab for experimental help and suggestions during the project. We are also indebted to Samantha Gruenheid and Lindsay Burns (McGill University, Montreal), John Brumell (SickKids, Toronto), Marc Vidal, David Hill, and Michael Calderwood (DFCI, Boston), Fritz Roth (University of Pittsburgh), and Robert Kozak (Sunnybrook Research Institute, Toronto) for sharing effector ORF plasmids or pathogen DNA for the construction of the eORFeome. We also thank Alexei Savchenko and Deepak Patel (University of Calgary) for their advice on Legionella RavE and Victor Manon and Jue Chen (Rockefeller University) for discussions on the TAP complex.

We thank Cassandra Wong of the Network Biology Collaborative Centre Proteomics Facility at the Lunenfeld-Tanenbaum Research Institute for mass spectrometry analysis. The facility is supported by the Canada Foundation for Innovation and the Ontario Government. Tomas Pachano was supported by the EMBO (906-2022) and VIP-2 Postdoctoral Fellowships, He Leng by a CIHR Postdoctoral Fellowship (FRN 187814), Guillaume Dugied by the EPIC Convergence Postdoctoral Fellowship Award, and Yeojin Lee by the CIHR REDI fellowship (#ED6-190718). Research in the Stark group is supported by Boehringer Ingelheim GmbH, the Austrian Research Promotion Agency (FFG, FO999902549), the Austrian Science Fund (10.55776/P36971, 10.55776/PAT3564423) and the WWTF (10.47379/LS24012). For the purpose of Open Access, the author has applied a CC-BY public copyright license to any Author Accepted Manuscript version arising from this submission. The Taipale group was supported by a Pathway Grant by the Temerty Faculty of Medicine of the University of Toronto and the Canada Research Chairs Program. The Schramek group was supported by a Terry Fox Research Institute Program Projects Grant (TFRI Project #1107), a Project Grant from the Canadian Institute of Health Research (CIHR PJT-496918) and the Canada Research Chairs Program. The Weirauch and Kottayan groups were supported by National Institutes of Health (NIH) R01 HG010730, R01 NS099068, R01 AI024717, and R01 AI148276.

## STAR* METHODS

### Key RESOURCES TABLE

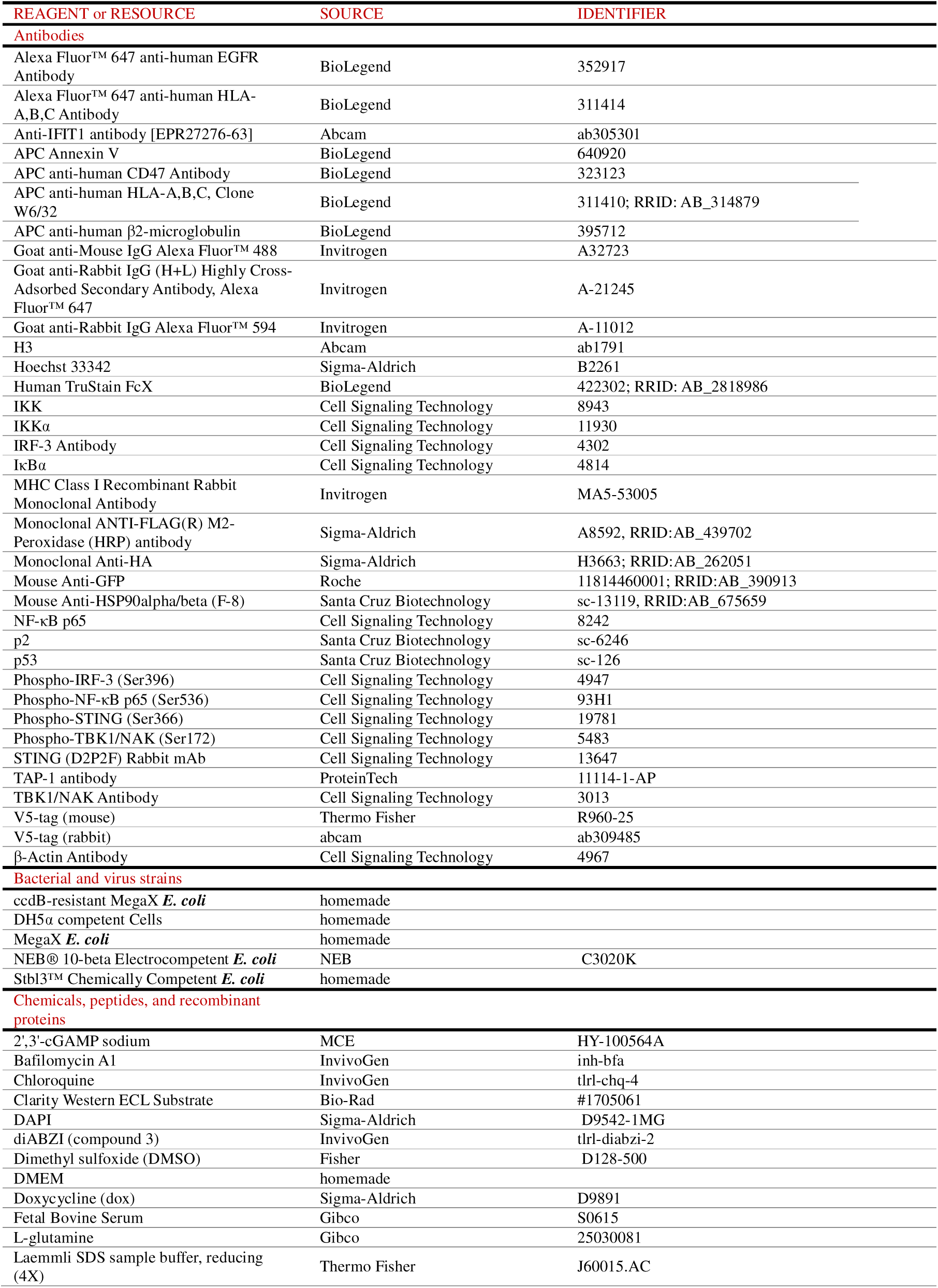
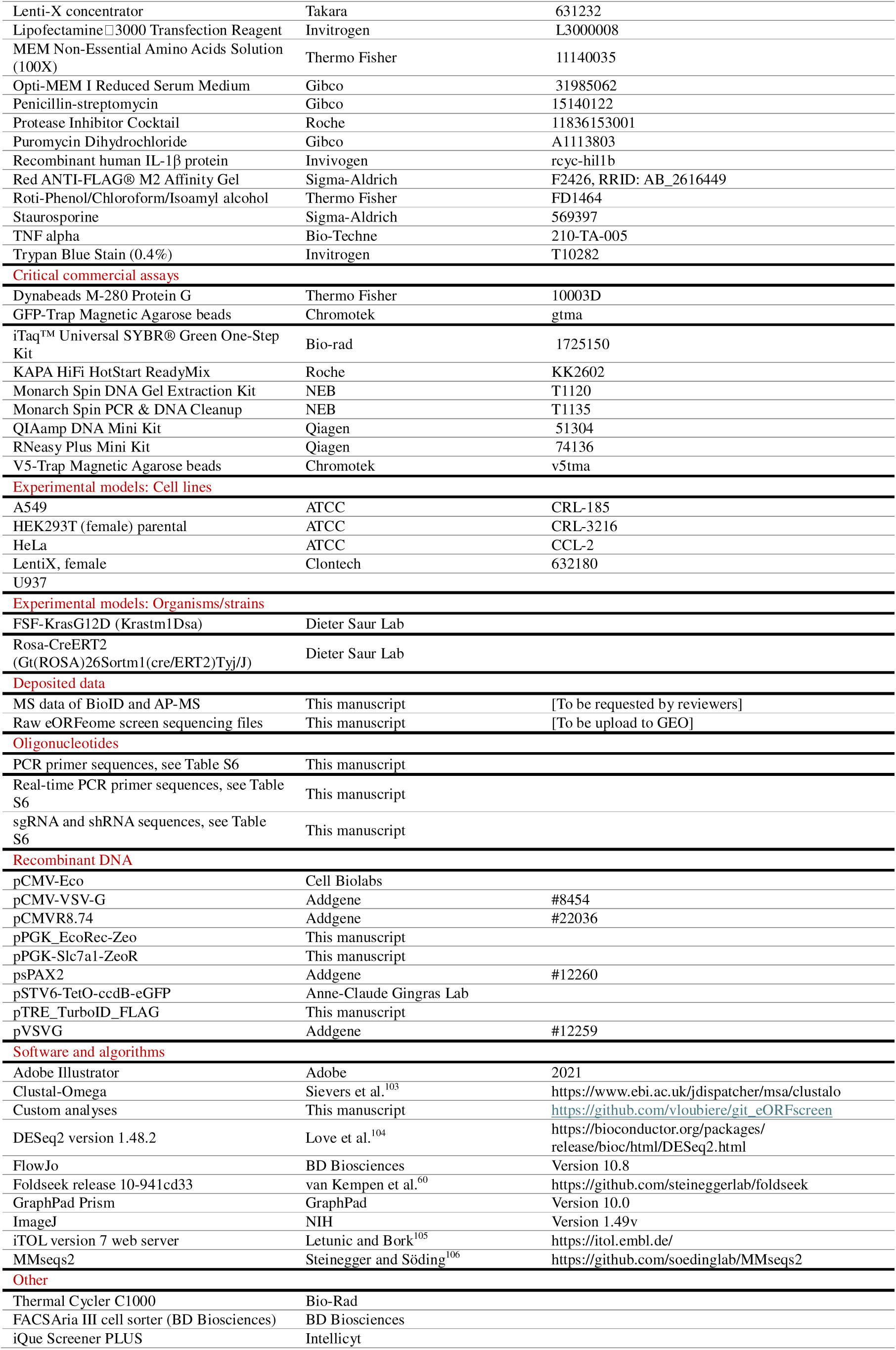

## Materials availability

All plasmids and cell lines generated in this study are available from the authors upon request.

### Data and code availability

All raw sequencing will be deposited in the Gene Expression Omnibus (GEO). Mass spectrometry datasets has been deposited with the ProteomeXchange Consortium via the MassIVE repository (https://massive.ucsd.edu/ProteoSAFe/static/massive.jsp) under identifier PXD069484 (BioID, token: 8AlWP6z4zZNH01iC), PXD069483 (FLAP-AP, token: 4jp66PQSf7IXJQAZ), PXD069487 (total proteome, token: jRlUb68t8CZ0610b). The datasets will be made public upon acceptance of the manuscript. The code used in this study can be found at https://github.com/vloubiere/git_eORFscreen. Any additional information required to reanalyse the data reported in this paper is available upon reasonable request.

## EXPERIMENTAL MODEL AND SUBJECT DETAILS

### Cell Lines

A549 cells (ATCC, CRL-185), HeLa (ATCC, CCL-2), HEK293T (female, ATCC CRL-3216) and Lenti-293T lentiviral packaging cells (female, Clontech, 632180) were cultured in Dulbecco’s modified Eagle medium (Sigma-Aldrich) supplemented with 10% fetal bovine serum, 2 mM L-glutamine (Sigma-Aldrich) and 1× penicillin-streptomycin (Sigma-Aldrich).

U937 cells (gift from Jason Moffat lab) were cultured in RPMI supplemented with 10% FBS.

Cells were passaged every 2 days using TrypLE Express (Thermo Fisher, 12605010) for dissociation and maintained at <70% confluency. All cells were cultured at 37 °C in a humidified incubator with 5% CO_2_. All the cell lines were determined negative for mycoplasma. Cells were used for experiments within 10 passages from thawing.

## Animals

Animal husbandry, ethical handling of mice, and all animal work were carried out according to the guidelines approved by the Canadian Council on Animal Care and under protocols approved by the Centre for Phenogenomics Animal Care Committee (18-0272H). The animals used in this study were Rosa-CreERT2 (Gt(ROSA)26Sortm1(cre/ERT2)Tyj/J, in C57/Bl6 background)^79^ and FSF-KrasG12D (Krastm1Dsa, in FVBN background),^78^ kindly provided by Dr. Dieter Saur at Technische Universität München, Munich, Germany. Genotyping was performed by PCR using genomic DNA prepared from mouse ear punches. For CreERT2-injected mice experiments, intragastric injections of approximately 75 mg tamoxifen/kg body weight (a standard dose of 50 μL tamoxifen/corn oil solution) were administered on postnatal days (P) 4 and 5.

## METHOD DETAILS

### Virus production

Lenti-X cells were co-transfected with lentiviral plasmids, pCMVR8.74 helper (Addgene plasmid no. 22036) and pCMV-VSV-G (Addgene plasmid no. 8454) or pCMV-Eco (Cell Biolabs) envelope plasmids using polyethylenimine (PEI) transfection (MW 25,000, Polysciences) as previously described.^107^ Virus containing supernatant was clarified by centrifugation and filtered through a 0.45 µm filter. Target cells were infected at limiting dilutions in the presence of 4 μg/mL of polybrene.

### Generation of A549 Cells Expressing the Ecotropic Receptor

To enable screening of the Lenti-eORFeome library in human cells under Biosafety Level-1 (BSL-1) conditions, we engineered the A549 cell line to express the mouse-specific ecotropic receptor, Slc7a1. This strategy renders the cells permissive to mouse ecotropic-pseudotyped lentivirus while ensuring that the viral particles cannot infect receptor-negative human cells, which minimizes risks of vector instability and the generation of replication-competent lentivirus.^108,109^ To achieve this, the lentiviral plasmid pTP36_EcoRec-Zeo was constructed to express Slc7a1 under the control of the PGK promoter, with a downstream internal ribosome entry site (IRES) driving a Zeocin resistance gene. A549 cells were then transduced with VSV-g pseudotyped lentivirus produced from this plasmid. Following selection with Zeocin, a clonal cell line, designated A549-Eco, was established by sorting single cells into 96-well plates using a FACSAria III cell sorter (BD Biosciences).

### Generation of clonal reporter cell lines

To generate reporter cell lines for NF-κB and p53 signalling, two lentiviral constructs were created. The NF-κB reporter construct contained four tandem copies of an NF-κB response element driving a minimal promoter and a destabilized enhanced green fluorescent protein (eGFP-mODC(d2)). Similarly, the p53 reporter construct contained 13 tandem copies of a p53 RE driving a minimal promoter and mCherry. Both plasmids included a PGK promoter driving a neomycin resistance gene for selection.

Ecotropic pseudotyped lentivirus was produced for each construct and used to transduce A549-Eco cells. Following selection with G418, populations were FACS-purified by sorting single cells with low basal fluorescence (eGFP or mCherry) into 96-well plates to establish clonal lines. Clones were subsequently validated for reporter functionality. The A549-Eco-NF-κB-eGFP clones were treated with 20 ng/mL TNFα, and the clone with the highest eGFP induction was selected. Likewise, A549-Eco-p53-mCherry clones were treated with 2.5 µM Nutlin, and the clone exhibiting the strongest mCherry induction was chosen for subsequent experiments.

### eORFeome library

Effector open reading frames as Gateway-compatible entry clones were sourced from previously published pathogen-specific ORF collections^35–38,110,111^ or ordered from DNASU Plasmid Repository (Tempe, AZ). Additional clones were from cDNA collections of viral genes^112,113^ that were amplified with PCR and cloned into pDONR221 Gateway entry vector. Salmonella effector ORFs were amplified from Salmonella genomic DNA, provided by John Brumell (SickKids, Toronto). E. coli and C. rodentium effectors were amplified from genomic DNA provided by Lindsay Burns and Samantha Gruenheid (McGill University, Montreal). Toxoplasma effectors were cloned by RT-PCR from Toxoplasma total RNA provided by Sebastian Lourido (Whitehead Institute, Cambridge, MA).

### Generation of a *Coxiella burnetii* Gateway library

A library of *Coxiella burnetii* effector sequences was cloned into pDONR221 as previously described^35–37^. Briefly, 147 sequences were selected from the C. burnetii Nine Mile Phase II RSA439 genome based on their homology to eukaryotic domains, known effectors, or direct observation of translocation.^114–116^ Each ORF was PCR amplified using primers with flanking attB sites, with each product including an in-frame ATG-start site and an amber stop codon (F: ggggacaagtttgtacaaaaaagcaggcttcATG…; R: ggggaccactttgtacaagaaagctgggtcATC…). Each was then cloned into the pDONR221-ccdB vector (Invitrogen) using Gateway BP clonase II (Invitrogen) per manufacturer’s instructions and sequence verified.

### Generation of barcoded backbone

We developed the pCW57.3-TRE3G destination vector from pCW57.1 (Addgene #41393) by replacing the TET promoter with a TRE3G promoter to minimize leakage expression of pathogen effectors in our library.

The library backbone was prepared by digesting the pCW57.3 plasmid with AgeI-HF and SalI-HF (NEB). The digestion products were separated by agarose gel electrophoresis, and the resulting vector backbone and insert template fragments were excised and purified using the Monarch DNA Gel Extraction Kit (NEB). The vector backbone was subjected to two additional rounds of cleanup using the Monarch PCR & DNA Cleanup Kit (NEB).

The insert template was barcoded using a two-step PCR strategy. First, a 50-cycle linear PCR was performed with a forward primer containing a semi-random 30-bp barcode (NNNNNWSWSWSWSWSNNNNNWSWSWSWSWS, where N is any base, S is G or C, and W is A or T; see Table S6 for primer details). This reaction used KAPA HiFi HotStart ReadyMix (Roche) and was followed by a single fill-in cycle with a reverse primer to generate double-stranded DNA. The barcoded insert was then amplified for four cycles using primers that added the necessary overhangs for Gibson assembly. PCR products were purified at each step, first using an Exo-SAP (NEB) treatment followed by the Monarch PCR & DNA Cleanup Kit, and finally with AMPure XP beads (Beckman Coulter).

The barcoded insert was assembled into the prepared vector backbone using an in-house Gibson Assembly Master Mix at a 5-fold molar excess of insert to vector. The reaction was incubated at 50°C for 1 h, and the final assembled library was purified. The library was then transformed into electrocompetent ccdB-resistant MegaX E. coli (homemade) via electroporation. After recovery in SOC medium, the transformants were pooled and expanded in LB medium containing ampicillin and chloramphenicol. Finally, the bacterial culture was harvested, and the plasmid library was purified using a Plasmid Plus Giga/Mega Kit (Qiagen).

### Gateway cloning of eORFs into barcoded backbone

Entry clones from the eORFeome collection were organized into 18 standardized subpools, each containing ∼384 eORFs, and subcloned into the barcoded pCW57.3 (see above). LR recombination reactions were assembled in duplicate using 150 ng of each entry ORF subpool and 150 ng of destination vector with 1 µL Gateway LR Clonase II in a total volume of 5 µL. Reactions were incubated overnight at 25°C in TE buffer. On each of the subsequent two days, an additional 150 ng of destination vector and 1 µL of LR enzyme were added to each reaction, and the volume was adjusted to 5 µL with TE buffer.

LR products were electroporated into NEB 10-beta electrocompetent cells and plated on LB agar containing carbenicillin (100 µg/mL). Plates were incubated overnight at 30°C. Colony counts confirmed a coverage of ∼200 fold for each eORF. Colonies were pooled, resuspended in SOC on ice, pelleted, and plasmid DNA was prepared using an endotoxin-free midiprep kit. For downstream lentivirus production, structural and non-structural effectors were maintained as two separate libraries (eORFeome-L1 and eORFeome-L2).

### Illumina sequencing of the eORF-barcode dictionary

The eORFeome plasmid library (1 µg) was first linearized by digestion with I-SceI (NEB) and subsequently purified using the Monarch PCR & DNA Cleanup Kit (NEB).

To assemble the Tn5 transposome, equimolar amounts of Tn5ME-fw and Tn5MErev oligonucleotides were annealed. The resulting adapters were then incubated with an in-house Tn5 transposase at room temperature for 30 min. The tagmentation reaction was performed by incubating 12 ng of the linearized plasmid library with the assembled Tn5 transposome in a 2x tagmentation buffer at 55°C for 8 min. The reaction was immediately stopped and purified using AMPure XP beads (Beckman Coulter).

The tagmented DNA was size-selected by separating the products on a 2% agarose gel and excising fragments larger than 150 bp. DNA was extracted using the Monarch DNA Gel Extraction Kit. This size-selected library was then used as a template for PCR amplification using KAPA HiFi HotStart ReadyMix (Roche) with a primer pair where one primer was biotinylated.

The biotinylated PCR products were captured on M280 streptavidin beads (Thermo Fisher Scientific). The bead-bound DNA was then used as a template for a final PCR step to add Illumina sequencing adapters. The final library was subjected to two consecutive rounds of gel extraction, isolating fragments between 150 and 300 bp. The library was given a final purification with the Monarch PCR & DNA Cleanup Kit before being submitted for illumina paired-end sequencing.

### Screens

#### Pooled eORFeome screens

The pooled-eORFeome library was generated by packaging into ecotropic pseudotyped lentivirus through the transfection of Lenti-X cells with PEI. Viral supernatant was subsequently cleared of cellular debris by filtration through a 0.45-μm PES filter. Target cells, including A549-Eco-NF-κB-eGFP, A549-Eco-p53-mCherry, A549-Eco, or U937-Eco lines, were then transduced at a multiplicity of infection (MOI) below 0.2 to ensure single-copy integration while maintaining a library representation of at least 10,000-fold. Twenty-four h post-transduction, cells were selected with 1 µg/mL puromycin until a non-transduced control plate of cells was eliminated, a process that typically required 4–5 days, during which cells were washed twice daily with 1× PBS and received fresh medium. Following selection, eORF expression was induced with 1 µg/mL dox for 24 h.

Screen-specific protocols were then implemented. For the reporter-based screens, the NF-κB activation screen involved collecting a baseline sample of 15 million cells as the input control before the remaining cells were sorted for the top 5% of the GFP-positive population using a BD FACSAria III sorter. Conversely, for the inhibition screens, cells were first treated with either 20 ng/mL TNFα (NF-κB inhibition) or 2.5 µM Nutlin (p53 inhibition) for 16 h. After this treatment, a 15 million cell input sample was collected, and the remaining cells were sorted to isolate the GFP-low or mCherry-low populations (bottom 5%), respectively. For the cGAS-STING inhibition screen, transduced U937 cells were treated with 10 µg/mL cGAMP, resuspended in PBS with 2% FBS containing FVD-780 viability dye (1:1000), and incubated at 4°C for 30 min in the dark. After two washes, the pellet was fixed using the BD Cytofix/Cytoperm™ Fixation Buffer, followed by incubation at 4°C for 20 min. Cells were then washed twice with 1x BD Permeabilization Buffer and incubated overnight at 4°C with an anti-IFIT1 primary antibody (abcam ,ab305301, 1:500). The following day, cells were washed twice and incubated with an Alexa Fluor™ 647-conjugated goat anti-rabbit secondary antibody (Invitrogen, A-21245; 1:300) for 30 min at 4°C. After two final washes, cells were resuspended in FACS sorting buffer for analysis. In the MHC-I inhibition screen, transduced A549 cells were first incubated with human TruStain FcX (BioLegend, 422302; 1:200) for 10 min at room temperature to block non-specific binding, followed by staining with an APC-conjugated anti-HLA-ABC antibody (BioLegend, 311410; 1:200) for 40 min at 4°C. Following three washes in FACS buffer, the cells were subjected to FACS. For the apoptosis inhibition screen, transduced A549-Eco cells were split post-induction into a control group maintained in dox and a treatment group cultured with both dox and 0.1 µM etoposide. These populations were maintained for 10 days and were split at a 1:4 ratio upon reaching confluency. Similarly, for the cGAMP survival screen, transduced U937 cells were initially treated with 10 µg/mL cGAMP, with the concentration increasing by 1 µg/mL every two days over a 10-day period. To select for strong suppressors, the screen concluded with the addition of 50 µg/mL cGAMP for 2 days. Finally, for every screen, genomic DNA (gDNA) was isolated from all collected cell populations (input, sorted or treated) for subsequent enrichment analysis. All screens were performed in at least two independent biological replicates.

#### DNA Isolation, Barcode PCR/eORFeome PCR, and Sequencing

gDNA was isolated by lysing cell pellets overnight at 55°C in a buffer containing 10 mM Tris-HCl (pH 8.0), 5 mM EDTA, 100 mM NaCl, 1% SDS, and 0.5 mg/mL proteinase K. The lysate was then treated with RNase A (Qiagen) for 2 h at 37°C. Subsequently, gDNA was purified through two sequential extractions with phenol:chloroform:isoamyl alcohol followed by a final extraction with chloroform:isoamyl alcohol.

Sequencing libraries were prepared using one of two strategies, dictated by whether the eORF sequence itself or its associated barcode was amplified for quantification. For the MHC-I and cGAS-STING inhibitor screens, the eORF sequences were directly amplified from gDNA and processed as previously described.^117,118^ For all other screens, eORF-associated barcodes were recovered from gDNA. To this end, the entire gDNA yield from sorted/selected samples and 40 µg of gDNA from input samples were used as templates. To maintain library complexity, 40 individual PCR reactions were performed for each sample. Amplification was carried out for 25 cycles (65°C annealing) using KAPA HiFi HotStart ReadyMix (Roche). The primers were designed to anneal to the Illumina adaptor sequences directly flanking the barcode region. To enable sample demultiplexing, these primers included Illumina i5 and idx index sequences as 5’ overhangs.

Following amplification, all PCR reactions for a given sample were pooled. The pooled amplicons were concentrated and purified using the Monarch PCR & DNA Cleanup Kit (NEB), using six columns for each input sample and one column for each sorted/selected sample. The final purified barcode amplicons were gel-extracted and submitted for next-generation sequencing.

#### Hit validations

To validate individual hits, candidate eORFs were subcloned from entry vectors into the barcoded pCW57.3 (for hits from the NF-κB or p53 screens) or pSTV6-eORF-C-GFP-Puro lentiviral plasmid (for hits from the MHC-I or cGAS-STING screens) via Gateway cloning. Lentivirus for each candidate was produced (see above) and used to transduce the appropriate cells in the presence of 4 µg/mL Polybrene. eORF-expressing cell populations were selected with 1 µg/mL puromycin.

For functional validation, each candidate was tested in parallel with and without dox. To identify activators, cells were cultured for 24 h before analysis. To identify inhibitors, cells were cultured for 24 h (with or without dox) and then stimulated for 16 h with the relevant pathway agonist. In all validation experiments, cells expressing a control eORF (mCherry, eGFP or NanoLuc) were included as a negative control. Reporter fluorescence was quantified by flow cytometry using iQue Screener PLUS (Intellicyt).

#### RNA extraction and quantitative PCR

48 h post-induction, U937 cells expressing U21-HHV7 or NanoLuc, alongside uninduced control cells, were harvested. Total RNA was extracted using the RNeasy Mini Kit (Qiagen) according to the manufacturer’s protocol. Subsequently, 1 µg of total RNA was reverse transcribed using the iScript cDNA Synthesis Kit (Bio-Rad). Gene expression levels were normalized to the housekeeping gene *ACTB*. All primer sequences are listed in Table S5.

#### Western Blots

Cells were lysed directly in Laemmli buffer containing 10% β-mercaptoethanol and boiled at 98°C for 5 min. Protein lysates were resolved by SDS-PAGE on 4–15% Mini-PROTEAN TGX Precast Gels (Bio-Rad) and transferred to Immobilon-P PVDF membranes (Merck Millipore) using a wet-transfer system. Membranes were probed with the following primary antibodies: anti-V5-tag (Thermo Fisher, R960-25, 1:1,000), anti-IKKα (Cell Signaling Technology, 11930), anti-IKKβ (Cell Signaling Technology, #8943), anti-Phospho-NF-κB p65 (Ser536) (Cell Signaling Technology, 93H1), anti-IκBα (Cell Signaling Technology, 4814), anti-NF-κB p65 (Cell Signaling Technology, 8242), anti-p53 (Santa Cruz Biotechnology, sc-126), anti-p21 (Santa Cruz Biotechnology, sc-6246), anti-human EGFR (BioLegend, #323123), anti-MHC Class I (Invitrogen, MA5-53005), anti-STING (D2P2F) (Cell Signaling Technology, 13647), anti-Phospho-STING (Ser366) (Cell Signaling Technology, 19781), anti-TBK1/NAK (Cell Signaling Technology, 3013), anti-Phospho-TBK1/NAK (Ser172) (Cell Signaling Technology, 5483), anti-IRF-3 (Cell Signaling Technology, 4302), anti-Phospho-IRF-3 (Ser396) (Cell Signaling Technology, 4947), anti-GFP (Roche, 11814460001; RRID:AB_390913), anti-HA (Sigma-Aldrich, H3663; RRID:AB_262051), anti-H3 (Abcam, ab1791), anti-β-Actin (Cell Signaling Technology, 967) and anti-HSP90α/β (F-8) (Santa Cruz Biotechnology, sc-13119; RRID:AB_675659) and anti-MHC-I (Invitrogen, MA5-53005).

Detection was performed using HRP-conjugated secondary antibodies (anti-mouse HRP, 7076; anti-rabbit HRP, 074; Cell Signaling Technology, 1:10,000) and Clarity Western ECL Substrate (Bio-Rad). Blots were imaged using a ChemiDoc Imaging System running Image Lab software (Bio-Rad).

#### Immunofluorescence

For U14 experiments, cells expressing empty vector, U14-V5, or the U14 D124R-V5 mutant were seeded in 96-well plates. To induce the expression of the V5-tagged constructs, cells were incubated with dox for 24 h, followed by a 1 h treatment with 1.5 µM Nutlin. Following treatment, cells were seeded in 96-well plates, washed once with PBS, and fixed for 10 min at room temperature with 3.7% formaldehyde in PBS. Following fixation, cells were washed with cold PBS and permeabilized with 0.1% Triton X-100 in PBS for 10 min at room temperature. The cells were then washed three times with PBS. Non-specific binding was blocked by incubating the cells in 5% bovine serum albumin (BSA) in PBS for 30 min at room temperature. Subsequently, cells were incubated overnight at 4°C with primary antibodies diluted in 5% BSA/PBS. The primary antibodies used were mouse anti-p53 (Santa Cruz Biotechnology, sc-126) and rabbit anti-V5-tag (abcam, ab309485) for the detection of V5-tagged U14 or V5-tagged U14 D124R constructs. After primary antibody incubation, cells were washed three times with PBS containing 0.1% Tween-20. Cells were then incubated for 1 h at room temperature in darkness with secondary antibodies diluted 1:1000 in 1% BSA/PBS. The secondary antibodies were goat anti-mouse IgG conjugated to Alexa Fluor™ 488 and goat anti-rabbit IgG conjugated to Alexa Fluor™ 594. Following secondary antibody incubation, cells were washed three times with PBS. Nuclei were counterstained with DAPI for 5 min at room temperature. Finally, cells were rinsed with PBS before imaging. For quantification, the total number of cells for each cell type was determined by counting DAPI-stained nuclei. The number of cells containing p53 foci (Alexa Fluor™ 488) and V5 foci (Alexa Fluor™ 594) was then counted. The percentage of cells positive for each type of foci was calculated relative to the total cell count. This analysis was performed for three independent experiments, and the data are presented in graphs as the mean ± standard deviation (SD). For 13.6k experiments, HeLa cells stably expressing 13.6k-GFP were seeded at 8,000 cells per well (96-well format). The following day, expression of 13.6k was induced using dox. 24h after induction, cells were fixed with 4% paraformaldehyde for 15 min and washed with PBS three times. MCHI staining was performed using anti-MCHI fused with AF647 antibody (1:20) for 1h at RT. Cells were then washed with PBS three times and incubated with Hoescht (1mg/ml) in 2x SSC for 10 min prior to imaging. Imaging was performed using an Opera Parking Elmer Phenix automated confocal microscope. Images were taken using a 40X water objective.

#### Luminescent Cell Viability Assay

For cell viability assays, U937 cells were seeded into 96-well plates at a density of 2,000 cells per well. The cells were induced with 1 µg/mL dox for 24 h, followed by stimulation with the indicated STING agonists for an additional 24 h. To measure viability, 100 µL of CellTiter-Glo reagent (Promega) was added to the 100 µL of cell culture in each well. The plates were agitated on an orbital shaker for 5 min and then incubated for 5 min at room temperature to stabilize the signal. Luminescence was subsequently measured using a BioTek multimode microplate reader.

#### TAP transporter assay

HeLa cells stably expressing C-terminal eGFP-tagged 13.6k constructs were seeded (100,000 cells per well, 48-well format) in a total volume of 500 µL. The following day, expression of the effectors was induced by adding 1 µg/mL of dox. 48h after induction, the cells were washed with PBS, prior to semi-permeabilization with 0.25 mg/mL saponin in PBS 1x for 15 min. The semi-permeabilization buffer was removed and 100 µL of transport buffer solution of 10 mM ATP, 10 mM MgCl2 and 1 nM fluorescence peptide in PBS 1x was added for 15 min. The reaction was stopped by adding 900 µL of PBS 1x supplemented with 20 nM EDTA. The samples were directly analyzed by flow cytometry.

#### U14-V5 AP-MS

For U14 experiments, A549-Eco cells were transduced at a high MOI with lentivirus expressing either U14-V5 or an Empty-V5 control and were subsequently selected with 1 µg/mL puromycin. For each biological replicate (n=3), cells from a confluent 15 cm plate were washed twice with ice-cold PBS and harvested.

Cell pellets were lysed in IP lysis buffer (20 mM HEPES pH 7.3, 150 mM NaCl, 2 mM MgCl2, 0.25% NP-40, 10% glycerol, 0.3% Triton X-100) supplemented with cOmplete™ EDTA-free Protease Inhibitor Cocktail (Roche). Lysates were incubated on ice for 30 min and sonicated. The lysates were then clarified by centrifugation at 20,000 x g for 5 min at 4°C.

The cleared supernatants were incubated overnight at 4°C with rotation alongside V5-Trap Magnetic Agarose beads (Chromotek) that had been pre-washed with IP lysis buffer. The following day, the beads were washed three times for 10 min each with IP lysis buffer, followed by four 5-minute washes with a no-detergent buffer (20 mM Tris pH 7.5, 130 mM NaCl). The final washed beads were submitted to the Vienna BioCenter Core Facilities (VBCF) Proteomics facility for on-bead digestion and subsequent mass spectrometry analysis.

For 13.6k experiments, co-immunoprecipitation experiments were performed using HeLa stably expressing 13.6k-GFP constructs. 48h after induction with dox, the cells were washed with 1x PBS and lysed in NP40 lysis buffer (10mM Tris/HCl pH 7.5, 150 mM NaCl, 0.5 mM EDTA and 0.5% NP40) with protease inhibitors on ice for 30 min. The lysate was centrifuged at 15,000g for 15 min at 4°C. Proteins were immunoprecipitated using GFP-Trap beads (ProteinTech, gtma) at 4°C for 1 h. Beads were then washed 3 times using wash buffer (10mM Tris/HCl pH 7.5, 150 mM NaCl, 0.5 mM EDTA and 0.05% NP40) to remove non-specific interactions. The immunoprecipitated proteins were eluted from beads by adding loading buffer and heating at 95°C for 5 min. Samples were then loaded on 4%-12% Bis-Tris PAGE and analyzed by Western Blotting on nitrocellulose membranes. Membranes were blotted with anti-GFP (11814460001, Roche, 1:1000) and anti-TAP1 (11114-1-AP, ProteinTech, 1:1000).

#### U14 BioID

For the BioID proximity labelling experiment, A549-Eco cells were transduced at a high MOI with lentivirus expressing either U14-TurboID or an Empty-TurboID control and were subsequently selected with 1 µg/mL puromycin. For each biological replicate (n=3), cells from a confluent 15 cm plate were treated with 500 µM biotin for 10 min at 37°C. The cells were then washed twice with ice-cold PBS and harvested.

Cell pellets were lysed in RIPA buffer (50 mM Tris-HCl pH 7.5, 150 mM NaCl, 0.1% SDS, 0.5% sodium deoxycholate, 1% Triton X-100) supplemented with cOmplete™ EDTA-free Protease Inhibitor Cocktail (Roche) and 1 mM DTT. After a 1 h incubation at 4°C, lysates were clarified by centrifugation at 18,000 x g for 10 min.

The cleared supernatants were incubated with streptavidin magnetic beads (Dynabeads™ M-280 Streptavidin, Thermo Fisher) to capture biotinylated proteins. The beads were then subjected to a stringent series of washes: once with RIPA buffer, once with 2% SDS, once with Buffer 2 (50 mM HEPES pH 7.4, 500 mM NaCl, 1 mM EDTA, 0.1% sodium deoxycholate, 1% Triton X-100), once with Buffer 3 (10 mM Tris-HCl pH 7.5, 1 mM EDTA, 250 mM LiCl, 0.5% sodium deoxycholate, 1% NP-40), and finally, four times with a non-detergent wash buffer (20 mM Tris-HCl, 100 mM NaCl). After the final wash, the beads were flash-frozen in liquid nitrogen and submitted to the proteomics facility for on-bead digestion and mass spectrometry analysis.

#### Whole-cell proteomics and analysis by Tandem Mass Tagging-based proteomics

For whole proteome analysis, we collected 2 million U937 cells expressing either wild-type U21-HHV7, the truncation mutant U21-HHV7Δtail or RLuc. Cells were lysed with 5% SDS, 50mM TEAB. Lysates were sonicated for 3 cycles of 5sec on, 3 sec off at 30% amplitude using a Sonicator with 1/8” microtip. 25 µg of protein material was reduced at 20mM DTT for 10 min at 95C and alkylated with 40mM iodoacetamide for 30 min in the dark. Samples were brought up to final concentration of 5% SDS and phosphoric acid was added to a final concentration of 1.2%. 165 µl of S-Trap protein binding buffer (90% methanol, 100mM TEAB) was added to 27.5 µl of acidified lysate. Resulting mixture was passed through the micro column at 4000xg. The micro column was washed 4 times with the S-Trap protein binding buffer. Each sample was digested with 4 µg of trypsin (in 20 µl of 50mM TEAB) for 1hr at 47C. Prior to elution, 40 µl of 50mM TEAB pH 8 was added to the column. Peptides were eluted by centrifugation at 4000xg. Peptides were eluted 2 more times with 40 µl 0.2% formic acid and 40 µl of 50% acetonitrile+0.2% formic acid. Eluted peptides were dried down and stored at -40LC. For data-dependent acquisition (DDA) LC-MS/MS, 250ng protein equivalent of digested peptides were analyzed using a nano-HPLC (High-performance liquid chromatography) coupled to MS. The sample was loaded onto Evotip Pure per manufacturer instructions. Peptides were eluted from the column (cat#: EV-1137, 15cmx150 µm with 1.5 µm beads) with the 30SPD pre-formed acetonitrile gradient generated by an Evosep One system and analyzed on a timsTOF Pro 2. The Evosep was coupled to the timsTOF Pro 2 using a 10 µm diameter emitter tip. The column toaster was set to 40C. The total DIA protocol is 44 min. The MS1 scan had a mass range of 100-1700Da in dia-PASEF. TIMS settings were accumulation and ramp time of 100ms, and within the mobility range (1/K0) of 0.6 to 1.6V·s/cm2. Cycle time 2s. For MS2, 1 mobility window with 2 ramps was used for 32 mass windows, 29.8 Da wide with 5Da mass overlap. The mobility range was from 0.61/K0 to 1.451/K0. This was at a duty cycle of 100% and a ramp rate of 9.52Hz. 1+ ions are excluded from fragmentation using a polygonal filter. The auto calibration was off.

#### 13.6K and U21 UltraID and AP-MS

HAdV 13.6K and HHV7 U21 clones were transferred from the pDONR221 entry vector into a destination vector carrying a C-terminal UltraID-FLAG tag via Gateway recombination. Stable HeLa cell lines expressing the effector-UltraID-FLAG fusion proteins were then generated. Cells were grown to 70% confluence in 15 cm dishes before inducing gene expression with 1 µg/mL dox for 24 h. 50 µM biotin was then added to each plate for 30 min. Cell pellets were collected for downstream UltraID and AP-MS applications.

For biotinylation of UltraID analysis, the cells were treated with 50 µM biotin (Sigma, B4501) for 1 hr, rinsed, and harvested and frozen in PBS buffer. UltraID (n=2) samples were prepared for MS analysis by the NBCC Proteomics Core at the Lunenfeld-Tanenbaum Research Institute (Toronto, Canada). Frozen cell pellets were lysed in 1:4 (cell pellet weight:lysis buffer) using RIPA-lysis buffer (50mM Tris-HCl (pH 7.5), 150mM NaCl, 0.1% (w/v) SDS, 1% NP-40, 1mM MgCl2, 1mM EGTA, 0.5mM EDTA, 0.4% (w/v) sodium deoxycholate, 1mM PMSF, and 1x Sigma protease inhibitors). The lysate was sonicated (3 x 5sec, 2 sec off) at 30% amplitude using a 1/8” microtip. 250 units TurboNuclease and 10 µg RNase was added to each sample and incubated, with rotation, at 4LC for 30 min. Additional SDS was added to bring the final concentration to 0.4% SDS. Each sample was mixed well and centrifuged at 14,000rpm for 20 min at 4LC. 200 µl lysate was applied to 20 µl of pre-washed 50% slurry streptavidin beads (Cytiva 17-5113-01), rendered trypsin-resistant, and incubated at 4LC, rotating, for 3 h. After incubation, supernatant was removed and beads were moved to a new tube in RIPA-wash buffer (50mM Tris-HCl pH 7.4, 150mM NaCl, 1mM EDTA, 1% NP-40, 0.1% (w/v) SDS, 0.4% (w/v) sodium deoxycholate). Beads were washed one time with 2% SDS, 50mM Tris-HCl pH 7.5, two times with RIPA-wash buffer, one time with TNNE-wash buffer (25mM Tris-HCl pH 7.4, 150mM NaCl, 0.1% NP-40, 1mM EDTA), and three times with 50mM ammonium bicarbonate. Proteins on beads were digested with 0.5 µg of trypsin in 50 µl of 50mM ammonium bicarbonate, overnight at 37LC. 0.25 µg trypsin was added to the peptides and incubated at 37LC for 3 h. Peptides were moved to a new tube, beads were washed with 50 µl water and this wash was combined into the same tube. Formic acid was added to a final concentration of 5%. The solution was centrifuged at 14,000rpm for 5 min, 80% of the solution was moved to a new tube. Peptides were dried down and stored at -40LC until mass spectrometry analysis.

For AP-MS, frozen cell pellets were lysed in 1:4 (cell pellet weight:lysis buffer) using lysis buffer (50mM HEPES-NaOH pH 8.0, 100mM KCl, 2mM EDTA, 0.1% NP-40, 10% glycerol, 1mM DTT, 1mM PMSF, and 1x Sigma protease inhibitors). The lysate was sonicated (3 x 5sec, 2 sec off) at 30% amplitude using a 1/8” microtip. 250 units TurboNuclease and 10 µg RNase was added to each sample and incubated, with rotation, at 4LC for 15 min. Each sample was mixed well and centrifuged at 14,000rpm for 20 min at 4LC. 400 µl lysate was applied to 25 µl of pre-washed 50% slurry anti-FLAG M2 magnetic beads (Sigma-Aldrich M8823) and incubated at 4LC, rotating, for 3 h. After incubation, supernatant was removed and beads were moved to a new tube in lysis buffer, with no PMSF, DTT, or protease inhibitors. Beads were washed one time with FLAG rinsing buffer (20mM Tris-HCl pH 8.0, 2mM CaCl_2_). The last wash was removed and proteins on beads were digested with 7.5 µl of trypsin (100 ng/µl in 20 mM Tris-HCl pH 8.0), overnight at 37LC. Peptides were moved to a new tube and 2.5 µl trypsin (same concentration) was added to the peptides and incubated at 37LC for 3 h. Formic acid was added to a final concentration of 5%. Peptides were dried down and stored at -40LC until mass spectrometry analysis.

#### Histology

For H&E staining, the mice were euthanized 9 weeks after the induction of U14 or eGFP. The lungs were inflated with intratracheal injection of 4% PFA and fixed overnight in 4% PFA. The fixed lungs were transferred to 70% ethanol, embedded in paraffin, and sectioned at 5 µm for H&E staining at the Histology core at University Health Network (UHN), Toronto, Canada. For quantification of tumor area, NDP.view.2 software was used.

## QUANTIFICATION AND STATISTICAL ANALYSIS

### Screen Data analysis

After initial quality checks using fastqc, raw sequencing reads amplified from integrated eORFs after phenotypic selection (screen) or not (input) were trimmed using Trim Galore (v0.6.10) in order to remove adapter sequences and filter out low-quality bases.^119^ Trimmed reads were then aligned to custom indexes that either directly contained eORF sequences (MHC-I and cGAS-STING screens) or a custom dictionary of non-ambiguous barcodes (all other screens) associated to each tested eORF (see Dictionary analysis) using Bowtie2 (v1.3.1).^120^ Reads with low mapping quality – that could not be unambiguously aligned to a unique ORF/Barcode – were discarded using SAMtools (v1.22.1).

Finally, reads associated to each ORF/Barcode were counted using the Rsamtools package (v3.21). Resulting matrices were used to identify hits that were significantly enriched after phenotypic selection (screen) compared to unselected/unsorted (input) samples.

For MHC-I and cGAS-STING screens, where integrated eORFs were directly sequenced, hits were identified using DESeq2 (v1.38.3),^104^ and only the eORFs with at least 10 reads in at least two of the compared conditions were considered. For all the other screens, were the abundance of eORFs was inferred from their associated BCs, MAGeCK (v0.5.9)^49^ was used, with the following parameters: --sort-criteria ‘pos’ --norm-method median --remove-zero none. Of note, only the BC/ORFs combinations with at least 3 reads across all screen/input sample pairs were considered, after adding a pseudocount of 1.

The complete scripts detailing the analysis pipeline described above are publicly available at the following GitHub repository: https://github.com/vloubiere/git_eORFscreen.

### Dictionary analysis

To link eORFs to their corresponding barcodes, we used the eORF-Barcode dictionary which was prepared and sequenced as described in the earlier ‘Illumina sequencing of the eORF-Barcode dictionary’ section. This library consists of DNA fragments where each eORF is positioned at the 5’ end and its associated barcode is at the 3’ end (fragment structure: Tn5_adaptor>eORF>AttB2>22nt_spacer>illuminaFwd>30nt_BC>illuminaRev. Of note, the Tn5 adaptor sequence was used for sequencing, not the illumina Fwd adapter which was only used for screening). Consequently, Read 1 captured the 3’ end of the eORF sequence, while Read 2 captured the barcode sequence, and were used to identify the BCs that were unambiguously assigned to each eORF sequence First, the 30nt barcode (BC) sequence was extracted from read 2 using cutadapt to trim the illuminaFwd and IlluminaRev adaptor sequences from the 5’ and the 3’ ends (5’ trimming parameters: -g GCTCTTCCGATCT; 3’ trimming parameters: -a AGATCGGAAGAGC -m 25 -M 30). Only barcode sequences that were between 25 and 30nt long and matched the expected sequence pattern ([GC][AT]){4}[GCAT]{5}([GC][AT]){5}[GCAT]{2}) were considered for downstream analyses.

To extract the eORF sequence, three strategies were used: (i) read1 were trimmed using cutadapt (5’ trimming parameters: -g AAAAAAGTTGGCA; 3’ trimming parameters: -a GCCCAACTTTCTT -m 25) or 2 (ii) read 1 were trimmed using Trim Galore^119^ (parameters: --hardtrim5 30) or (iii) read 2 were trimmed using trim_galore (parameters: --hardtrim3 30). eORF sequences were then aligned to the sequences present in the eORFeome library using bowtie2 (v1.3.1, default parameters)^120^ and, for each read ID, only the best alignment (based on mapping quality was retained), and only the reads that were unambiguously aligned (mapping quality ≥ 30) were eventually retained.

Then, confident BC/eORF pairs were retrieved using the read ID, and only the BCs that were supported by at least three read counts and were systematically assigned to the same unique eORF were considered. These barcode sequences were finally used to generate the custom Bowtie2 index used to align the sequencing results from the phenotypic screens.

The complete scripts detailing this analysis is publicly available at the following GitHub repository: https://github.com/vloubiere/git_eORFscreen.

### Phylogenetic distribution of eORFs

Each eORF was first assigned the taxonomic identifier (TaxID) of its source pathogen from the NCBI Taxonomy database (https://www.ncbi.nlm.nih.gov/taxonomy). These TaxIDs were then used as input to generate a phylogenetic tree based on the NCBI taxonomic hierarchy, which was exported in Newick format. The resulting tree was visualized using the Interactive Tree Of Life (iTOL) web server.^105^

### Mass Spectrometry Analysis

For data-dependent acquisition (DDA) LC-MS/MS, one-sixteenth of digested peptides were analyzed using a nano-HPLC (High-performance liquid chromatography) coupled to MS. The sample was loaded onto Evotip Pure per manufacturer instructions. Peptides were eluted from the Performance column (cat#: EV-1109, 8cmx150 µm with 1.5 µm beads), heated at 40C) with the 60SPD pre-formed acetonitrile gradient generated by an Evosep One system, and analyzed on a timsTOF Pro 2. The Evosep was coupled to timsTOF Pro 2 using a 20um diameter emitter tip. The column toaster was set to 40C. The total DDA protocol is 22 min. The MS1 scan had a mass range of 100-1700Da in PASEF mode. TIMS settings were accumulation and ramp time of 100ms (with 4 PASEF ramps and active exclusion at 0.4min), and within the mobility range (1/K0) of 0.85 to 1.3V·s/cm2. This was at a cycle time of 0.53s. The target intensity was set to 17,500 and intensity threshold set to 1750. 1+ ions are excluded from fragmentation using a polygonal filter. The auto calibration was off.

For Whole-cell proteomics analysis, MS Data Analysis with Spectronaut, Spectronaut v19 directDIA+ workflow was used to search the data with the Spectronaut generated Human spectral library (Human_PDB_2023). Parameters for the search were default. Differential abundance testing used was unpaired t-test.

For UltraID and AP-MS, Mass spectrometry data generated were stored, searched and analyzed using ProHits laboratory information management system (LIMS) platform. Within ProHits, MSFragger 4.1 was used to search the data using a FASTA database from UP000005640 human Uniprot proteome, no isoforms, with a custom list of contaminants and decoy appended. Acetylated protein N-term and oxidated methionine were set as a variable modifications. Precursor mass tolerance was set to 20ppm on either side. Fragment mass tolerance was set to 20ppm. Enzymatic cleavage was set to trypsin with 2 missed cleavages. MSBooster and Percolator were turned on. Percolator required a minimum probability of 0.5 and did not remove redundant peptides. The target-decoy competition method was used to assign q-values and PEPs. For ProteinProphet, the maximum peptide mass difference was set to 30ppm. When generating the final report, the protein FDR filter was set to 0.01. FDR was estimated by using both filtered PSM and protein lists. Razor peptides were used for protein FDR scoring. All other parameters were default.

### Homology clusters

To group eORFs by sequence similarity, protein sequences were clustered using MMseqs2 v13.45111^106^ across a range of identity thresholds (0.8, 0.5, 0.3, 0.15 and 0.1).

### UniProt annotation scores

UniProt annotation scores for eORFs were retrieved from UniProt using their assigned UniProt, CRC64 or RefSeq IDs. In case it matches with more than one entry, the highest score was kept. To note, some viral entries may map to the precursor polyprotein rather than the specific eORF segment. eORFs lacking an annotation score indicate no exact sequence match to any UniProt entry. The human proteome analysis was restricted to reviewed Homo sapiens entries (organism ID: 9606; reviewed:true; n = 20,420).

### Statistical analysis

GraphPad Prism v10 and RStudio with R v4.2.0 were used for all statistical analysis. Statistical tests are described in figure legends.

### Clustering of eORF hits

For the clustering, only eORFs significantly enriched in any of the barcoded analyzed with MAGeCK^49^ (FDR ≤ 0.05 and LFC ≥ log2(1.5)) or non-barcoded screens analyzed with DESeq2^104^ (FDR ≤ 0.01 and LFC ≥ 1) were considered. Log fold change (LFC) values were clipped at the 10th and 90th percentiles and LFC of non-significant hits was set to zero. Hits were then clustered using a two-layer self-organizing map using the supersom function from the kohonen R package (v3.0.10) with a 4×5 hexagonal, toroidal grid. The first layer contained the clipped LFC values (user.weight = 1), and the second layer used a binary matrix indicating significant hits (user.weight = 10).

### Enrichment of homologs

For each cluster of eORF hits defined in the previous section, significantly over-represented protein homology groups (defined as having ≥30% sequence homology using MMseqs2,^106^ as detailed above) were identified using a one-sided Fisher’s exact test (alternative= ‘greater’), with all the other clusters constituting the control group. Of note, only protein homology groups containing ≥ 2 ORFs were considered. Resulting p-values were adjusted for multiple hypothesis testing across all cluster-homology group comparisons using the Benjamini-Hochberg method to control the false discovery rate (FDR).

### Structural modelling

Predicted three-dimensional structures of the U14-p53 complex and of monomeric RavE were generated using the AlphaFold 3 server (https://alphafoldserver.com/).^121^ The UniProt accession numbers Q69549 (U14) and P04637 (p53) were provided as input for the complex prediction, while Q5ZZ16 was used for RavE. PDB database (https://www.rcsb.org/) was used to retrieve the models of DNA-p53 (PDB ID: 2AC0), SV40 LargeT-p53 (PDB ID:2H1L) and HPV16 E6-p53 (PDB ID: 8GCR). All the resulting structural models were visualized using UCSF ChimeraX (v1.10).^122^

To identify structural homologs, the predicted structure of RavE (amino acids 1-234; UniProt: Q5ZZ16) was used as a query to search the PDB100 database (release 20240101) using Foldseek.^60^ The search was taxonomically restricted to Bacteria.

---

**Supplementary Figure 1.**
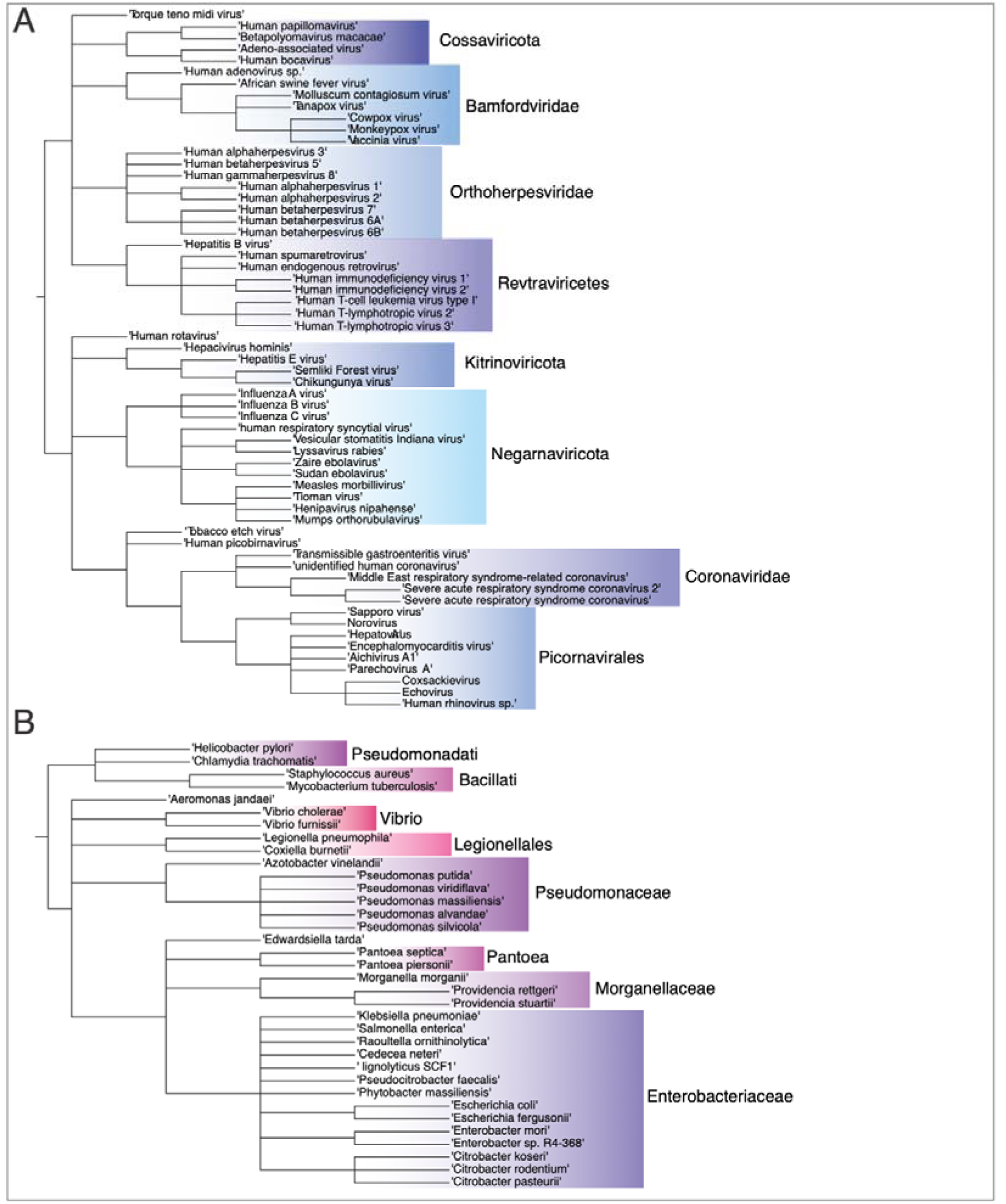
Evolutionary relationships of eORFs in the eORFeome library. Phylogenetic tree of the viral (A) or bacterial (B) eORFs present in the eORFeome library.

**Supplementary Figure 2.**
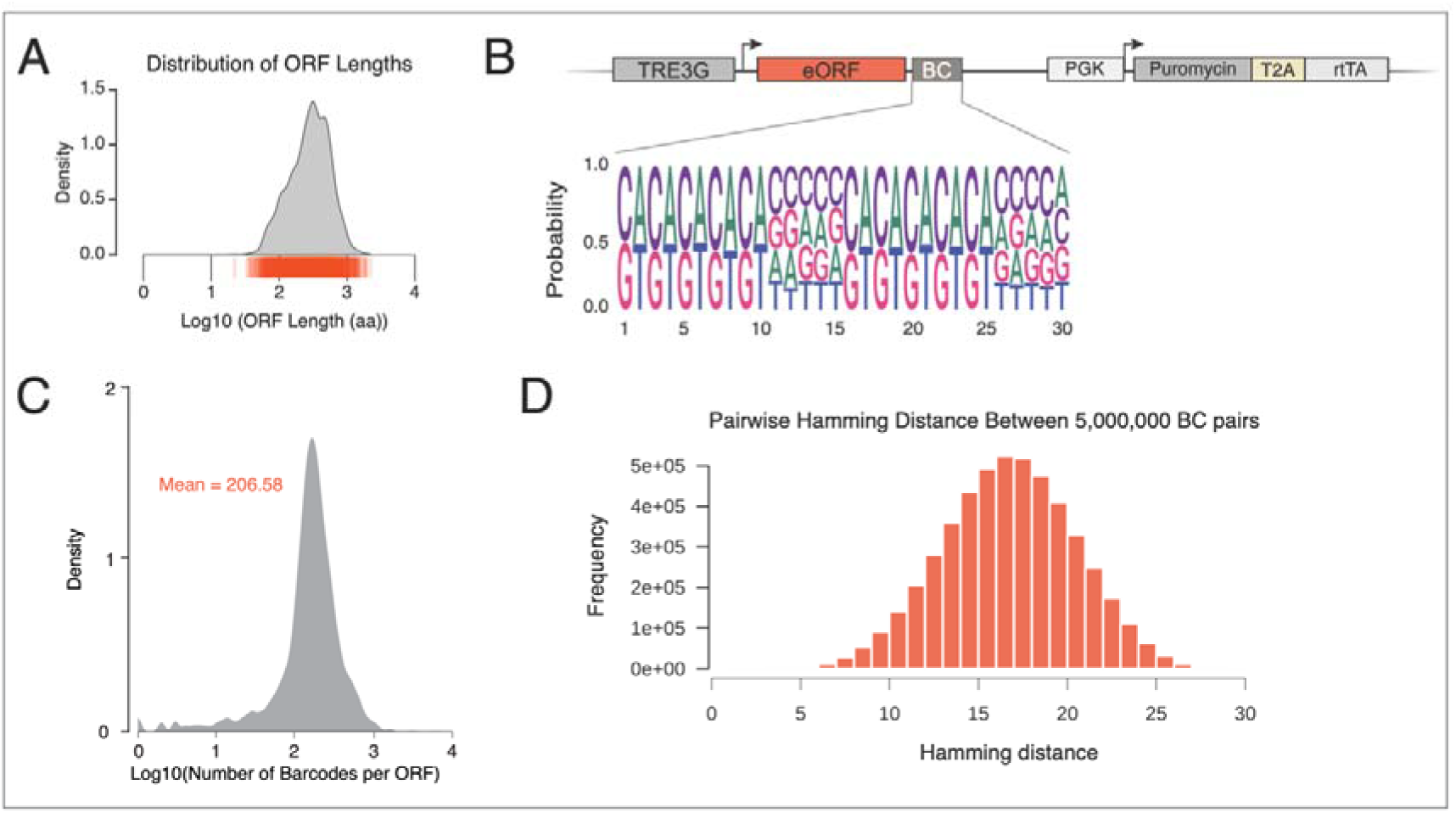
Characterization of the eORFeome library. (A) Distribution of eORF lengths. The density plot is based on the Log10-transformed length (in amino acids) of each ORF in the library. The underlying orange lines marks the position of each individual eORF. (B) Top: Scheme of the lentiviral construct, detailing the arrangement of its key components: a dox-inducible TRE3G promoter drives the expression of an eORF, which is followed by a unique barcode (BC). Downstream of this, a constitutive promoter ensures the expression of both puromycin for cell selection and rtTA, which is essential for the dox-inducible activation of the TRE3G promoter. Bottom: position weight matrix that describes the semi-random nature of the 30-nucleotide barcode (BC), which consists of the pattern [(SW)×5 + N×5] repeated twice, where S represents G or C, W represents A or T, and N represents any nucleotide. (C) Density plot of the Log10 distribution of the number of barcodes per ORF, with a mean of 206.58 BC per ORF. (D) Histogram of the Hamming distance distribution, where the frequency (y-axis) of a given Hamming distance (x-axis) was calculated from 5 million randomly sampled barcode pairs from the library.

**Supplementary Figure 3.**
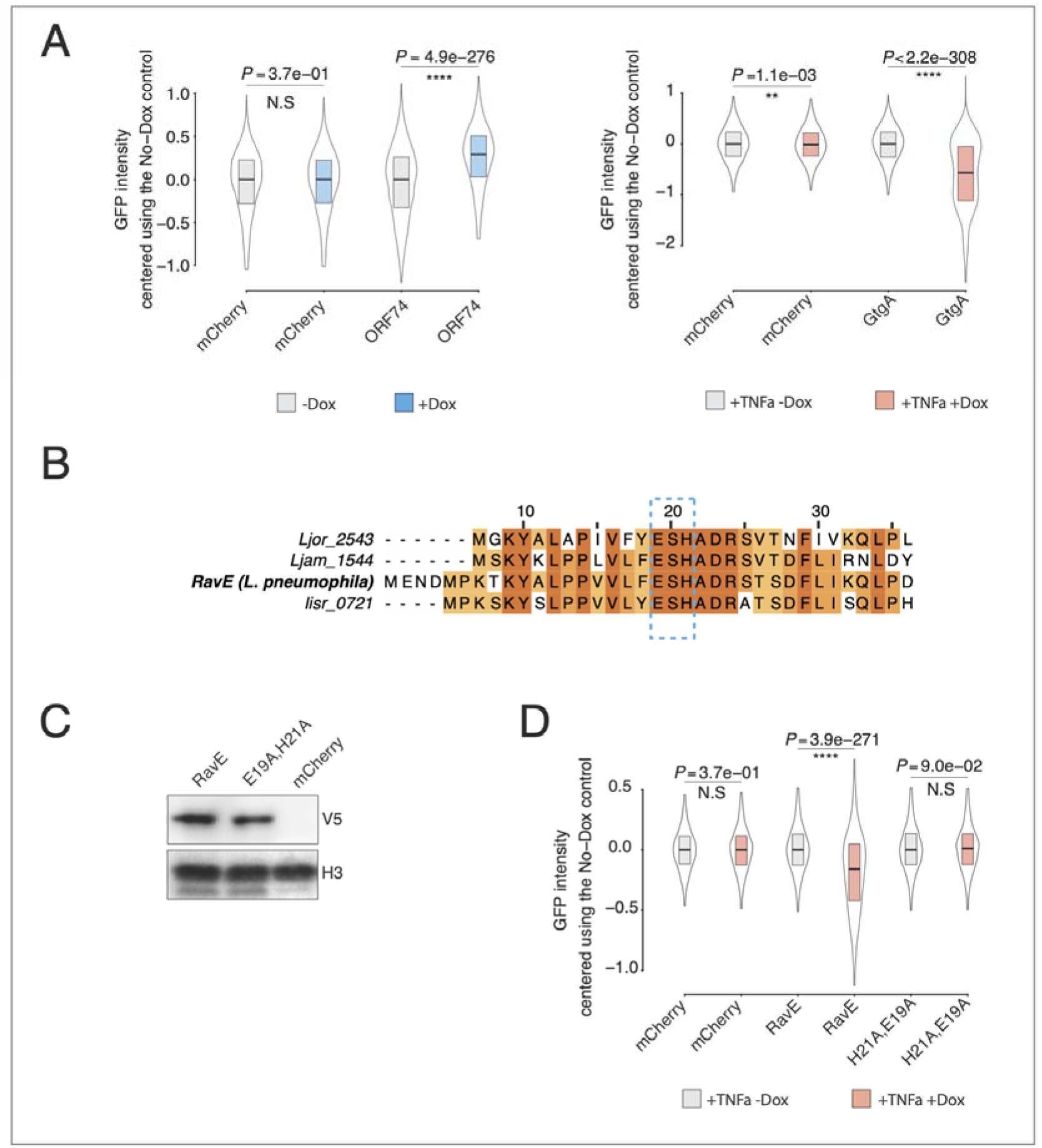
**Validation of eORF hits from NF-**κ**B Screens.** (A) Boxplots showing representative examples of the single-cell GFP fluorescence distribution, complementing the median-based analysis presented in Figure 2B. ORF expression was induced with dox (blue or orange) or left untreated (grey) and where specified, cells were treated with 20 ng/mL TNFα for 16 hours to stimulate NF-κB activity. The y-axis represents the Logicle-transformed GFP fluorescence, centred to the median of the corresponding no-dox control population. Boxplots display the median and interquartile range. mCherry serves as the negative control. Statistical significance was determined by comparing dox-treated versus untreated samples for each ORF. (B) Multiple sequence alignment of four RavE homologs. Conserved residues are coloured in orange. (C) Western blot analysis of cells expressing V5-tagged RavE, V5-tagged RavE E19A,H21A mutant, or an mCherry control. Lysates were immunoblotted with an anti-V5 antibody to confirm that the mutant protein is stably expressed. H3 served as a loading control. (D) Boxplots showing single-cell GFP fluorescence distributions, generated using the same analysis method described in (A). These data complement the median-based analysis presented in Figure 2J.

**Supplementary Figure 4.**
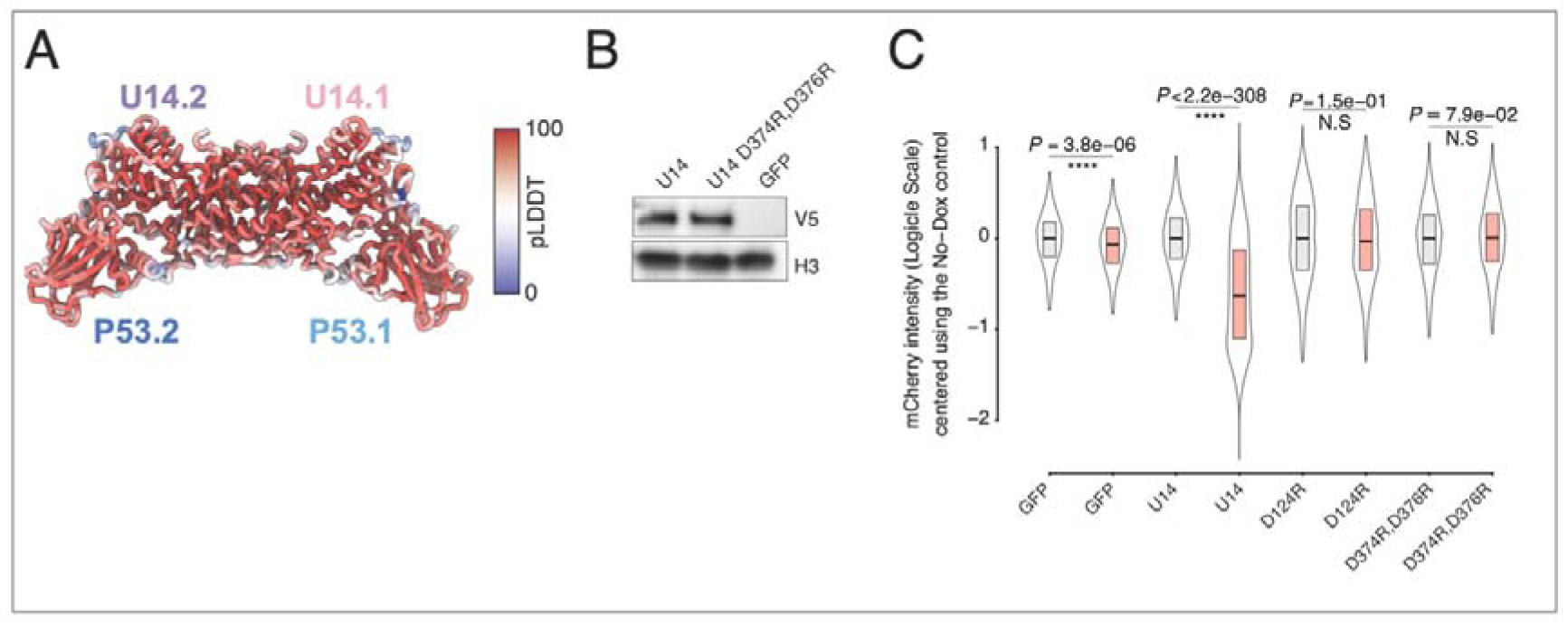
Validation of U14 from HHV6A as a p53 antagonist. (A) Multiple sequence alignment of four RavE homologs. Conserved residues are coloured in orange. (B) Western blot analysis of cells expressing V5-tagged U14, V5-tagged U14 D124R mutant, or an GFP control. Lysates were immunoblotted with an anti-V5 antibody to confirm that the mutant protein is stably expressed. H3 served as a loading control. (C) Predicted local distance difference test (pLDDT) scores for the U14–p53 complex structural model generated by AlphaFold 3. (B) Western blot analysis of cells expressing V5-tagged U14, V5-tagged U14 D374R,D376R mutant, or an mCherry control. Lysates were immunoblotted with an anti-V5 antibody to confirm that the mutant protein is stably expressed. H3 served as a loading control. (C) Boxplots showing representative examples of the single-cell mCherry fluorescence distribution, complementing the median-based analysis presented in Figure 4. ORF expression was induced with dox (orange) or left untreated (grey) and where specified, cells were treated with 2.5uM nutlin for 16 hours to stimulate p53 activity. The panels show results for wild-type U14 with its inactive mutants (D124R or D374R,D376R). The y-axis represents the Logicle-transformed mCherry fluorescence, centred to the median of the corresponding no-dox control population. Boxplots display the median and interquartile range. GFP serves as the negative control. Statistical significance was determined by comparing dox-treated versus untreated samples for each ORF. *padj < 0.05, **padj < 0.01, ***padj < 0.001, **** padj < 0.0001.

**Supplementary Figure 5.**
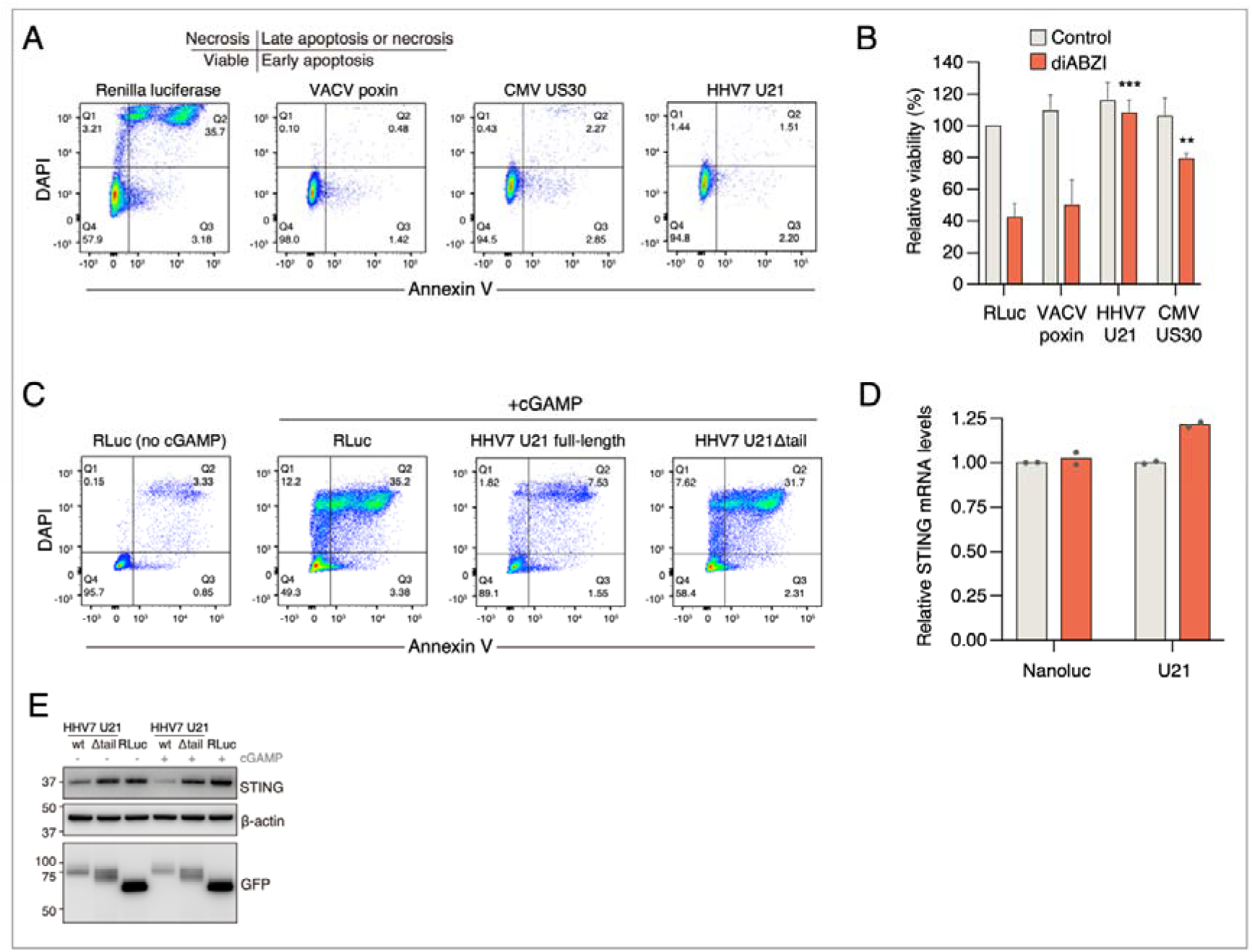
Validation of U21 as a potent inhibitor of STING signaling. (A) U937 cells stably expressing dox-inducible GFP-tagged Renilla luciferase, VACV poxin, CMV US30, or HHV7 U21 were induced with dox followed by 2’,3’-cGAMP treatment and analyzed for apoptosis with Annexin V and DAPI staining. (B) The same constructs were analyzed for viability CellTiter-Glo after treating the cells with the non-nucleotide STING agonist diABZI. (C) Cells expressing full-length HHV7 U21 or U21Δtail were analyzed for apoptosis after cGAMP treatment as in (A). (D) STING mRNA levels were assessed by qRT-PCR in cells expressing Nanoluc-GFP or U21-GFP. (E) STING protein levels were analyzed by western blotting in cells expressing the indicated GFP-tagged constructs after treating the cells with cGAMP or vehicle.

**Supplementary Figure 6.**
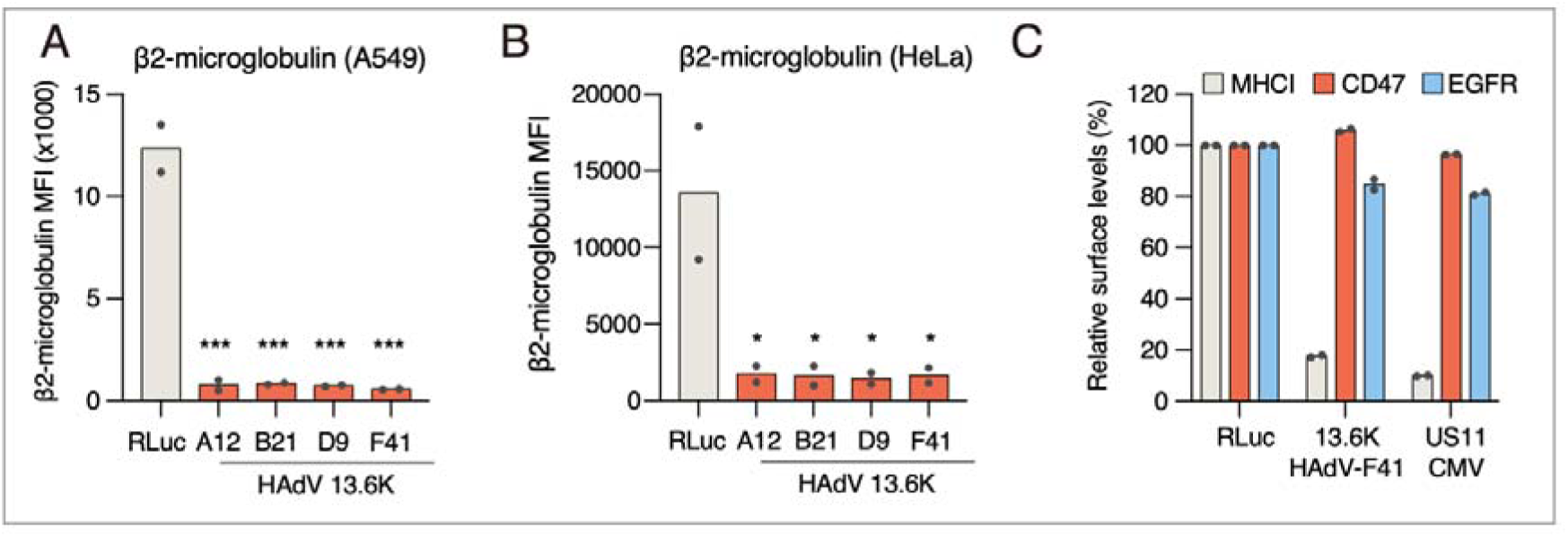
Validation of 13.6K proteins as novel TAP inhibitors. (A-B) A549 cells (A) and HeLa cells (B) expressing indicated GFP-tagged constructs were assessed for cell surface β2-microglobulin levels by flow cytometry. (C) HeLa cells expressing GFP-tagged HAdV-F41 13.6K or CMV US11, a known inhibitor of MHC-I surface display, were analyzed for cell surface MHC-I, EGFR, or CD47 levels by flow cytometry.

## Notes

### Competing Interest Statement

The authors have declared no competing interest.

